# Catalytic and inhibitory architecture of comammox ammonia monooxygenase

**DOI:** 10.64898/2026.09.02.748776

**Authors:** Tie-Qiang Mao, Xiaoyun Yang, Zhi-Cong He, Jiawen Yang, Rihui Wu, Kenneth M.Y. Leung, Shengying Li, Ping Han, Wei Peng, Zongqiang Li

## Abstract

Complete ammonia oxidizers (comammox) are widespread nitrifiers that can dominate ammonia oxidation in diverse environments by efficiently converting ammonia to nitrate within a single cell, yet the molecular basis distinguishing their ammonia monooxygenase (AMO) from those of canonical bacterial AMO remains unresolved. Here we report cryo-electron microscopy (cryo-EM) structures of AMO from the comammox *Nitrospira inopinata* (*Ni*AMO) in inhibitor-free and allylthiourea (ATU)-bound states at 2.47 Å and 2.68 Å resolution, respectively. *Ni*AMO displays distinctive auxiliary-subunit organization, copper-site configuration and hydrophobic-channel architecture. Integrative molecular dynamics (MD) and quantum mechanics/molecular mechanics (QM/MM) calculations support a methyl-plastoquinol (methyl-PQH_2_)-coupled, Cu_D_-centric catalytic model, with Cu_C_ potentially facilitating quinone redox cycling. *N. inopinata* exhibited broad susceptibility to several known nitrification inhibitors, and ATU-bound *Ni*AMO structure localized the inhibitor to the Cu_C_–Cu_D_ region, accompanied by constriction of the hydrophobic channel, which is consistent with the competitive role of ATU demonstrated in recovery assays. Multi-omics analyses further revealed an energy-limited stress response to ATU, including induction of urea transport and utilization systems. Collectively, these findings define a methyl-PQ-linked catalytic and inhibitor-responsive architecture of comammox AMO and establish a mechanistic framework for lineage-aware management of nitrification in natural and engineered ecosystems.

## Introduction

Nitrification is a cornerstone process in global nitrogen cycle and governs primary productivity, nitrogen retention and climate-relevant emissions^1,2^. Ammonia oxidation, the first and rate-limiting step of nitrification, is catalyzed by the integral membrane-bound ammonia monooxygenase (AMO), which converts NH_3_ to hydroxylamine. AMO provides the primary energy-conserving reaction for ammonia-oxidizing archaea (AOA), ammonia-oxidizing bacteria (AOB) and complete ammonia oxidizers (comammox)^1,3–6^. AOA and AOB convert ammonia to nitrite, whereas nitrite-oxidizing bacteria (NOB) subsequently oxidize nitrite to nitrate^7^.The discovery of comammox *Nitrospira*, which oxidize ammonia to nitrate within a single cell, overturns this long-standing division of nitrification between ammonia and nitrite oxidizers^3,6^. Comammox are now recognized across soils, freshwaters, sediments and engineered systems^8–11^. Their ecological reach extends to coastal Antarctica, where clade B comammox were abundant and actively incorporated ^13^CO_2_ during nitrification at low temperature^12^. The cultured clade A species *N. inopinata* also has exceptionally high ammonia affinity^13^ and can grow on guanidine as its sole source of energy, reductant and nitrogen; soil microcosms further suggest that guanidine-derived nitrogen can enter nitrification in agricultural systems^14^. These observations place comammox at the intersection of natural nitrogen cycling, extreme-environment microbiology and managed nitrogen inputs.

As an integral copper-dependent metalloenzyme, copper-site configuration and catalytic mechanism of AMO have long been central topics of investigation. Recent AOB AMO structures have begun to reveal lineage-dependent subunit composition and copper coordination, where the simultaneously occupied Cu_C_–Cu_D_ has been proposed to act concertedly with the reduced coenzyme Q (CoQ10H_2_) as the physiological electron donor, while a conformational rearrangement of TM5 in the AmoC subunit enlarges the hydrophobic substrate channel^15–18^. Whether comammox AMO shares these structural features or instead couples a distinct active-site organization and reductant coupling to reactive oxygen-species generation and ammonia oxidation remains unresolved.

Beyond its biogeochemical significance, ammonia oxidation contributes to nitrous oxide (N_2_O) production^19–22^ and accelerates the conversion of fertilizer ammonium to mobile nitrate, promoting nitrogen loss, water contamination and eutrophication^23–26^. To address these issues, nitrification inhibitors (NIs), including nitrapyrin (2-chloro-6-(trichloromethyl)pyridine, NP), dicyandiamide (DCD), amidinothiourea (ASU), 3,4-dimethylpyrazole phosphate (DMPP), and allylthiourea (ATU), are therefore used to inhibit AMO activity^27–29^. Their efficacy varies with soil properties and ammonia-oxidizer composition, and some compounds affect non-target organisms^27,30,31^. Comammox and canonical AOB can respond differently to the same inhibitor, while comammox responses also vary among soils^32,33^. Since comammox contribute to nitrification in agricultural systems and can access alternative nitrogen substrates such as guanidine^14,34^, inhibitors developed primarily around canonical AOB may not predictably control comammox activity. The molecular basis of this lineage-dependent inhibition remains unresolved because no inhibitor-bound structure of comammox AMO has been available.

Here, we report cryo-EM structures of the *Ni*AMO, captured in inhibitor-free and inhibitor-bound states. These structures reveal distinct structural and mechanistic features of its auxiliary subunit composition, regulation of the substrate access channel, and the functional specialization between the Cu_C_ and Cu_D_ active sites. Structural and computational analyses support an inclined mononuclear Cu_D_ active site that exhibits elevated catalytic energy barrier with a species-specific reductant methyl-PQH_2_, consistent with the lower catalytic activity of *Ni*AMO relative to AOB AMOs. Inhibition assays show that *N. inopinata* has the greatest susceptibility to most tested inhibitors among the three representative nitrifiers examined. Rescue assays suggest that ATU inhibits *Ni*AMO via substrate-competitive chelation of its intracellular copper centers. The ATU-bound structure directly visualizes a multilayered inhibitory mechanism involving copper chelation, channel closure and impaired reductant access. Multi-omics profiling further links direct ATU-mediated *Ni*AMO inhibition to an energy-limited cellular response and increased urea-acquisition capacity. Together, these findings link structural divergence, methyl-PQH_2_-coupled redox chemistry and inhibitor sensitivity in comammox AMO, providing a molecular framework for lineage-aware interpretation and management of nitrifiers.

## Results

### Structure of *N. inopinata* AMO

To determine the structure of comammox AMO, we cultivated *N. inopinata* and prepared an AMO-enriched membrane fraction (Supplementary Fig. 1). Subsequent cryo-EM single-particle analysis yielded a high-resolution reconstruction at a nominal resolution of 2.47 Å (Supplementary Fig. 2). Similar to previously characterized bacterial AMOs and pMMOs^15–17^, *Ni*AMO forms a trimeric assembly in which each protomer comprises the canonical subunits AmoA, AmoB and AmoC, along with a newly identified transmembrane helix (hereafter Helix1) (Fig. 1a, b and Supplementary Fig. 3a-d). In contrast to the auxiliary helices observed in bacterial AMOs and pMMOs, Helix1 resolved in *Ni*AMO occupies a distinct structural position that enables intimate association with both AmoA and AmoC through tightly bound lipid molecules, and adopts a tilted orientation of approximately 55° relative to the membrane-embedded equatorial plane (Fig. 1b and Supplementary Fig. 4). By integrating the cryo-EM density, mass spectrometric data and genomic annotation from *N. inopinata*^3,6^, we proposed the Helix1 as a structural fragment derived from the putative AmoD subunit or a downstream enzyme—such as nitrite oxidoreductase (NXR) (Supplementary Fig. 3c-e). Unexpectedly, three rod-like electron densities were found in the periplasmic cupredoxin domains of *Ni*AMO. These elongated densities are buried within a pocket at the base of the β-barrel formed predominantly by bulky side-chain (Fig. 1c, d and Supplementary Fig. 3f); their chemical identity and functional significance remain unknown.

**Fig. 1.**
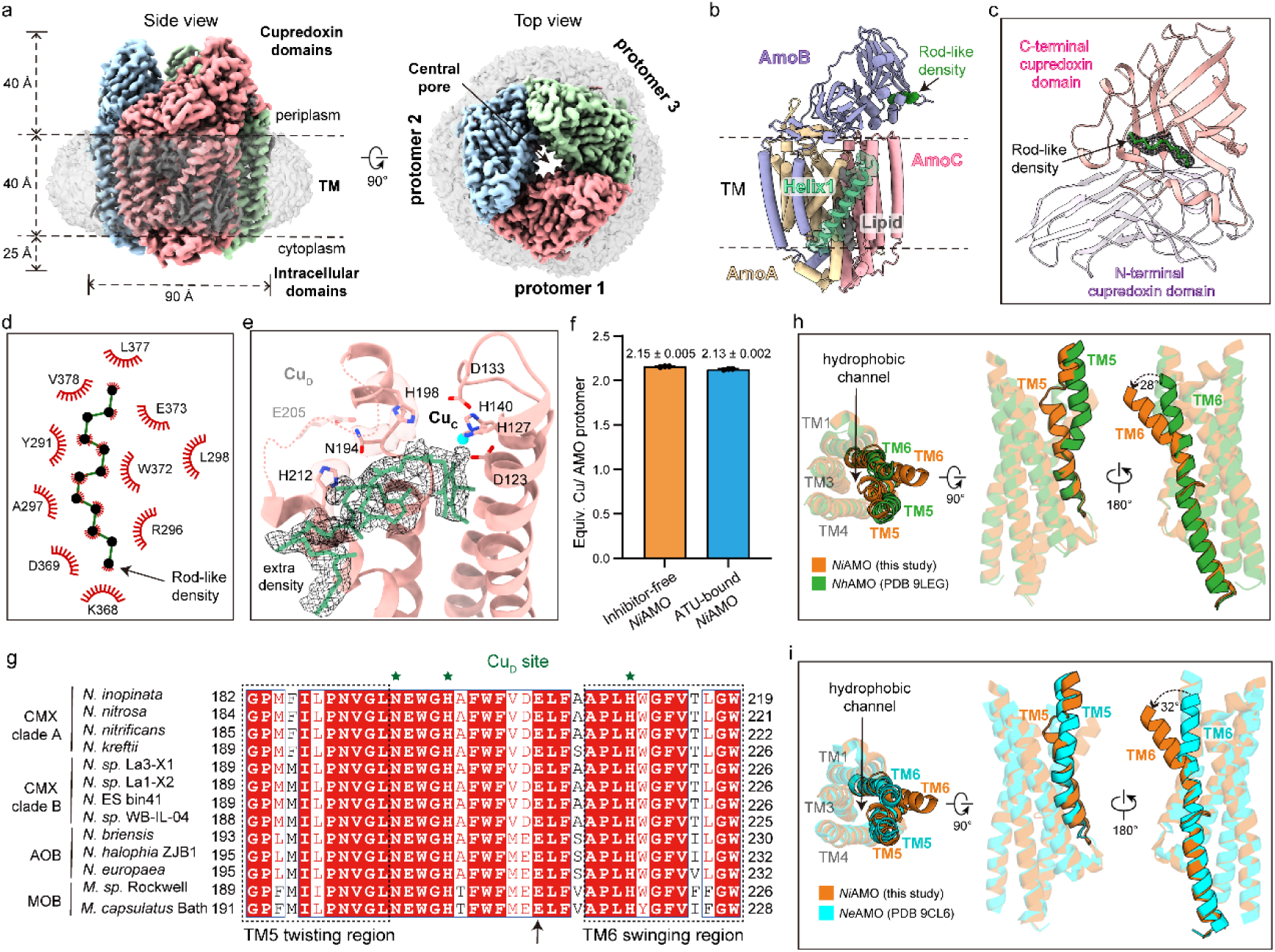
Overall structure of AMO from *N. inopinata.* **a,** Cryo-EM map of *Ni*AMO shown in two views. The three protomers are colored pink, blue and green; lipid densities are colored gray and the membrane bilayer is displayed as a transparent density. **b,** Model of one *Ni*AMO protomer, including AmoA (yellow), AmoB (purple), AmoC (pink), and Helix1 (green) with the corresponding density map. Lipid densities bridging Helix1 and the AMO core are depicted in gray. **c,** The rod-like density observed within the cupredoxin domains of AmoB. The N- and C-terminal cupredoxin domains are colored in purple and coral, and the unidentified density is shown in green. **d,** Close-up view showing the interactions between residues and the rod-like molecule. **e,** Coordination environments of Cu_C_ and Cu_D_ sites. An additional density is tentatively modelled as quinone-like molecule (green). The unresolved TM5-TM6 loop is depicted as dashed lines. **f**, Copper content of inhibitor-free *Ni*AMO and ATU-bound *Ni*AMO from untreated and ATU-treated *N. inopinata* cells. Error bars represent standard deviation of n = 3, each measured in triplicate. **g,** Sequence alignments from different AOMs and MOBs demonstrate that the residues located within the twisting TM5 region, TM6 swinging region and Cu_D_ coordinated residues are highly conserved. Red stars represent the Cu_D_ coordinated residues. Black arrow indicates the conserved E205 residue. **h-i,** Structural comparisons of hydrophobic channels in *Ni*AMO (colored in orange) with AOB AMOs containing co-occupied Cu_C_–Cu_D_ center (**h**, PDB: 9LEG, green) and a mononuclear copper center (**i**, PDB: 9CL6, cyan).

The high-resolution structure of *Ni*AMO enables unambiguous modeling of the Cu_B_ and Cu_C_ sites (Fig. 1e and Supplementary Fig. 3b, g-h). In contrast, the Cu_D_ site lacks well-defined electron density, attributable to the conformational flexibility of the TM5-TM6 loop (Fig. 1e). Recent biochemical and structural evidence indicate that the Cu_D_ site may constitute the catalytically active center of pMMO^35,36^ and a dicopper center comprising Cu_C_ and Cu_D_ is simultaneously present in AOB AMOs^15,17^. To quantify copper occupancy, we performed inductively coupled plasma (ICP) mass spectrometry analysis. The results showed that copper content in inhibitor-free *Ni*AMO is comparable to that in the inhibitor-bound state (Fig. 1f), in which three copper ions in per protomer of AMO supported by cryo-EM structure (*vide infra*), suggesting that the Cu_D_ site is occupied in *Ni*AMO. Owing to the conformational flexibility of the TM5-TM6 loop, the conserved Cu_D_-coordinating E205 remains unresolved, whereas the three other conserved Cu_D_-coordinating residues-H212, H198 and N194-are clearly resolved in TM5 and TM6 respectively (Fig. 1e, g), suggesting that the E205 is more susceptible to structural heterogeneity at the Cu_D_ site.

Consistent with the recently identified nonprotein density near Cu_C_ and Cu_D_ sites in bacterial AMO^15,17^, a quinone-like density was observed in the vicinity of this Cu_C_ and the unresolved Cu_D_ sites (Fig. 1e). Recent biochemical assays demonstrated that reduced coenzyme Q may function as a physiologically relevant electron donor for AMO activity of AOB^17^, whereas spectrometric and biosynthetic studies identified methyl-PQs in comammox *Nitrospira*^37,38^. The shape, dimensions and spatial orientation of the quinone-like density are compatible with a methyl-PQH_2_ scaffold. The modeled methyl-PQH_2_ headgroup is oriented toward both Cu_C_ and Cu_D_, suggesting an efficient short-range electron transfer to facilitate ammonia oxidation^39^. Together with the coenzyme Q-linked electron donation reported for AOB AMO, this observation suggests lineage-specific coupling between AMO and quinone chemistry. The transmembrane (TM) helices surrounding Cu_C_–Cu_D_ region also distinguish *Ni*AMO from the published AOB AMO structure. In canonical AOB AMO structures with co-occupied Cu_C_ and Cu_D_ sites, outward displacement and local kinking of TM5 from AmoC enlarge the hydrophobic channel^15,17^. By contrast, TM5 in *Ni*AMO adopts a slightly more inward orientation, resembling structures with a mononuclear copper center (Fig. 1h, i). The upper segment of TM6 instead rotates by 24°-32° toward the central pore, resulting in an enlarged entrance of the hydrophobic channel (Fig. 1h, i and Supplementary Fig. 5). These structural differences indicate that TM6, rather than TM5, serves as the primary regulatory element governing the Cu_D_ coordination and the hydrophobic channel geometry in *Ni*AMO. Both the upper segments of TM5 and TM6 are highly conserved across diverse bacterial lineages, including comammox (clades A and B), AOB and methane-oxidizing bacteria (MOB) (Fig. 1g), suggesting that lineage-specific use of a conserved helical module generates distinct channel states and copper-site configurations. When considered together with the quinone-like density adjacent to the Cu_C_–Cu_D_ region, these structural observations raise the possibility that methyl-PQH_2_ may influence not only local electron transfer but also the dynamic organization of the Cu_D_ site.

### Cu_D_ dynamics

Despite the comparable copper content of inhibitor-free and inhibitor-bound *Ni*AMO (Fig. 1f), the Cu_D_ site is poorly resolved in the inhibitor-free state. This lack of well-defined density coincides with pronounced conformational flexibility of the TM5–TM6 loop surrounding the Cu_D_ region (Fig. 1e), raising the possibility that Cu_D_ is dynamically coordinated rather than constitutively absent. To investigate whether the presence of methyl-PQ influences Cu_D_ coordination, we performed long-timescale MD simulations using inhibitor-free *Ni*AMO structure without (Supplementary Fig. 6) or with methyl-PQH_2_ (Supplementary Fig. 7). A free Cu ion was initially placed within the Cu_D_ coordination environment and a plausible geometry was generated based on the bacterial AMO model under distance and angle restraints^17^. The restraints were then released to evaluate the intrinsic dynamics of Cu_D_ site.

In the methyl-PQH_2_-free structure, interaction between Cu_D_ site and N194, H198 and H212 progressively weakened and the Cu ion moved away from the coordination region (Supplementary Fig. 6). By contrast, in the methyl-PQH_2_-bound *Ni*AMO model, these residues maintained a persistent coordination environment around the Cu ion throughout the simulations (Supplementary Fig. 7). These observations support a potential association between methyl-PQH_2_ and enhanced stability of the Cu_D_ coordination. However, compared with the Cu_C_ coordination, Cu_D_ exhibits substantially greater conformational flexibility (Supplementary Figs. 6c and 7d), suggesting the intrinsically dynamic nature of Cu_D_ site. Free-energy calculations further established a direct energetic link between methyl-PQH_2_ binding and stabilization of the Cu_D_ coordination environment. In methyl-PQH_2_-free *Ni*AMO, formation of the Cu_D_-coordinating state requires overcoming an energy barrier of 19.9 kcal/mol and is strongly endergonic by 15.8 kcal/mol relative to the non-coordinating state (Supplementary Fig. 8). By contrast, in methyl-PQH_2_-bound *Ni*AMO, the corresponding barrier is reduced to 7.7 kcal/mol, and the free-energy penalty of the Cu_D_-coordinating state decreases to 4.5 kcal/mol (Fig. 2a-d). These results indicate that methyl-PQH_2_ could facilitate the conformational reorganization required for Cu_D_ coordination and stabilize the resulting coordination environment. Nevertheless, the Cu_D_-coordinating state remains moderately endergonic (4.5 kcal/mol) (Fig. 2c, d), indicating that it is likely transiently or only partially populated rather than constitutively occupied. Such dynamic and incomplete Cu_D_ occupancy precludes unambiguous Cu_D_ density in the cryo-EM reconstruction, despite the preservation of three of the four Cu_D_-coordinating residues, consistent with the absence of Cu_D_ site in the recently reported AOB AMO structure^16^. Collectively, these data indicate that Cu_D_ undergoes dynamic conformational change, and methyl-PQH_2_ may facilitate transient assembly and stabilization of the Cu_D_ coordination environment.

**Figure 2.**
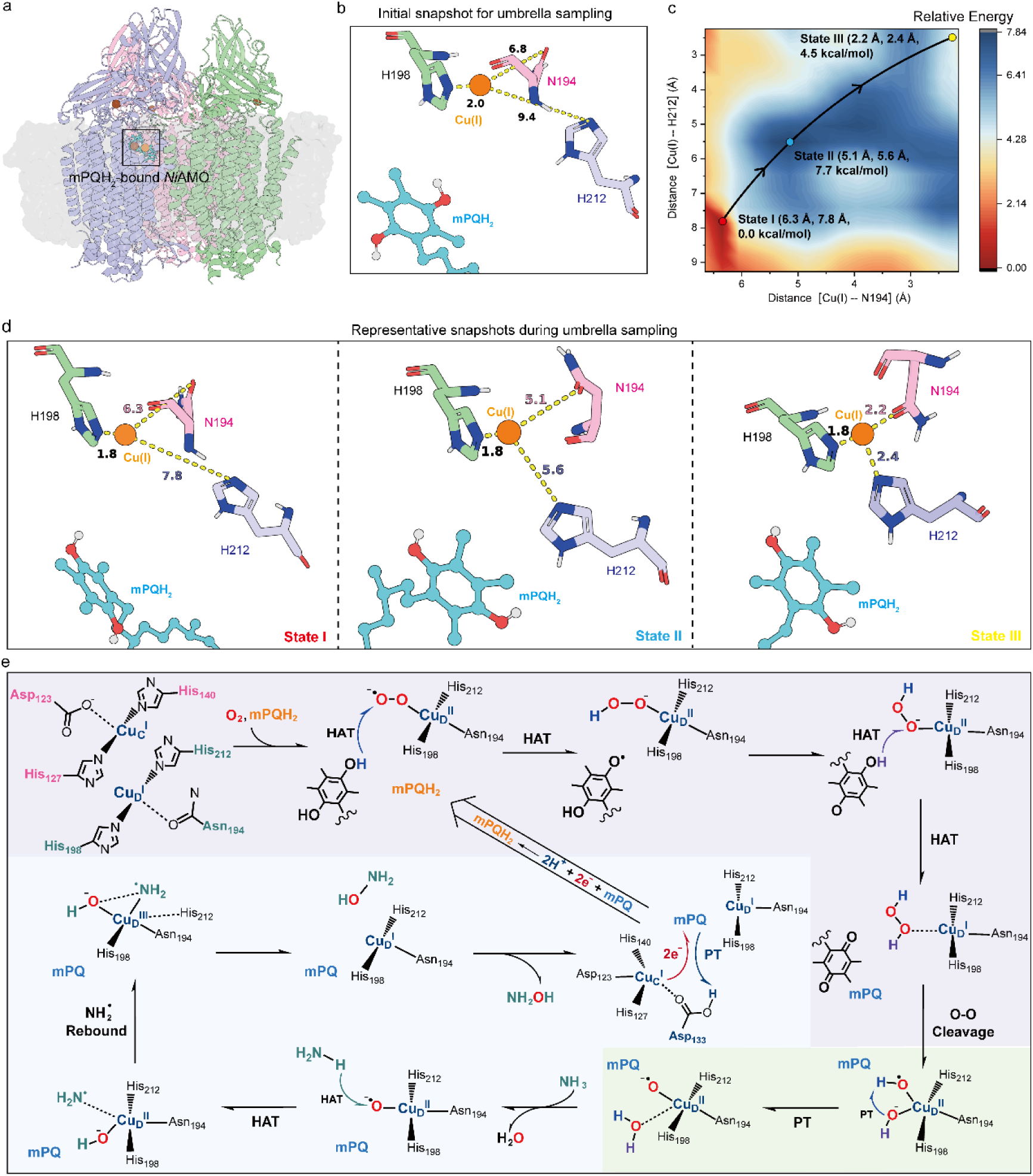
Mechanistic insights into methyl-PQH_2_-affected Cu_D_ binding and formation of the reactive copper-oxygen species. **a,** Overall structure of the membrane-bound *Ni*AMO-methyl-PQH_2_ complex. The membrane is shown as a gray surface. The Cu(I) ion of Cu_C_ site and methyl-PQH_2_ resolved in the cryo-EM structure are colored as brown and cyan, respectively, whereas the Cu(I) ion randomly placed near the Cu_D_ site are shown as orange spheres. **b,** Initial snapshot for umbrella sampling (US) simulations, in which Cu(I) was placed near H198. N194, H198, and H212 are depicted as pink, green, and blue stick models, respectively. **c,** US-calculated the free energy profile (in kcal/mol) for the transition of Cu(I) from the initial non-coordinating state (State I) to the Cu-coordinating state (State III). The reaction coordinates were defined as the distances between Cu(I) and N194 and between Cu(I) and H212, respectively. **d,** Representative snapshots sampled along the transition pathway of Cu(I) from the uncoordinated to the coordinated state. Key distances are given in angstrom (Å). **e,** Overview of the proposed catalytic cycle mediated by the Cu_D_ active center. The reaction is initiated by O_2_ activation at the Cu_D_(I) site (purple background), followed by the formation of Cu_D_-associated reactive oxygen species Cu_D_(II)-O^•–^ generation (green background), substrate oxidation steps, ultimately leading to product formation (blue background).

### Methyl-PQH_2_-mediated formation of the reactive copper-oxygen species

To further probe the role of methyl-PQH_2_ in *Ni*AMO catalysis, we simulated *Ni*AMO with methyl-PQH_2_ and simultaneously occupied Cu_C_ and Cu_D_ sites. During the trajectories, the Cu_C_–Cu_D_ distance progressively increased (∼ 15 Å), whereas the quinol headgroup of methyl-PQH_2_ preferentially approached the Cu_D_ center (∼ 6 Å, Supplementary Fig. 9). This geometry favors a catalytic configuration centered primarily at Cu_D_, distinguishing *Ni*AMO from the more tightly coupled Cu_C_–Cu_D_ model proposed for AMO of AOB^17^ (two copper distance is ∼ 8 Å ^18^).

QM/MM and QM/MM metadynamics calculations were subsequently used to delineate how methyl-PQH_2_ promotes O_2_ activation and hydroxylation of NH_3_ (Fig. 2e, Supplementary Tables 1-3). In the computed pathway, methyl-PQH_2_ donates two hydrogen-atom equivalents sequentially to the Cu_D_(II)–O_2_^•⁻^ species, resulting in the formation of a Cu_D_(I)···H_2_O_2_ intermediate (RC_CuD_ → IC3_CuD_, Supplementary Fig. 10). Following the first hydrogen atom transfer (HAT), the resulting methyl-PQH radical undergoes a pronounced conformational reorganization before the second transfer takes place (IC1_CuD_ → IC2_CuD_, Supplementary Fig. 11). The highest computed electronic-energy barrier for conversion of Cu_D_(II)–O_2_^•⁻^ to Cu_D_(I)···H_2_O_2_ is 14.6 kcal/mol (^3^RC_CuD_ → ^3^TS1_CuD_), and the *in situ* formation of H_2_O_2_ is exergonic by 19.8 kcal/mol (^1^IC3_CuD_). Notably, the initial HAT step mirrors that in AOB AMO, forming a Cu_D_(II)–OOH^⁻^ intermediate. However, the subsequent reactivity of this intermediate depends on the distinct Cu_C_–Cu_D_ geometry. In AOB AMO, the shorter Cu_C_–Cu_D_ distance enables Cu_D_(II)–OOH^⁻^ to interact with Cu_C_(II), yielding a bridging dicopper oxygen species (two copper distance is ∼ 8 Å ^18^). By contrast, the longer dicopper distance and greater steric constraint in *Ni*AMO prevent this bimetallic coupling, diverting the intermediate toward H_2_O_2_ formation and mononuclear Cu_D_-mediated O–O bond cleavage (Fig. 2e). This mechanism echoes a previously proposed mode of reactive oxygen-species generation in pMMO, in which a quinol cofactor participates directly in assembling the oxidizing species at a mononuclear copper center^39^.

The newly formed H_2_O_2_ is directly recruited for the subsequent catalytic step. Electron transfer (ET) from Cu_D_(I) weakens and cleaves its O–O bond with a barrier of 17.5 kcal mol⁻^1^, producing Cu_D_(II)–OH⁻ together with a hydroxyl radical (•OH) (IC3’_CuD_ → IC4_CuD_, Supplementary Fig. 12). A subsequent HAT, accompanied by H_2_O formation, converts this transient radical pair into the reactive Cu_D_(II)–O^•⁻^ species (Fig. 2e). Thus, methyl-PQH_2_ drives a confined process of O_2_ reduction, H_2_O_2_ formation and O–O bond cleavage that generates a copper–oxygen species capable of substrate oxidation. Once formed, Cu_D_(II)–O^•⁻^ activates an N–H bond of NH_3_ and promotes the ensuing formation of NH_2_OH with an overall barrier of 17.2 kcal mol⁻¹ (Fig. 2e and Supplementary Fig. 13). Collectively, the calculations support a Cu_D_-centered pathway for oxygen-activation and ammonia-hydroxylation.

The Cu_C_ site may instead facilitate local regeneration of the quinol reductant. In the modeled reaction, oxidized methyl-PQ is first converted to the methyl-PQH^•^ radical through a proton-transfer (PT)-triggered electron-transfer process, in which protonated D133 supplies the proton while Cu_C_(I) provides the electron. This first reduction step proceeds with a barrier of only 1.2 kcal mol⁻^1^ (Fig. 2e and Supplementary Fig. 14). The resulting methyl-PQH^•^ radical subsequently undergoes conformational reorientation, positioning it for a second proton-coupled reduction that generates methyl-PQH_2_ with a barrier of 9.1 kcal mol⁻^1^ and an overall reaction free energy of −9.3 kcal mol⁻^1^ (Supplementary Fig. 15). The calculations implicate D133, located adjacent to Cu_C_, as a plausible proton donor during this local redox process (Supplementary Fig. 16).

Notably, the intermediate Cu_C_–D133 distance observed in the cryo-EM structure may be compatible with interconversion between a protonated, non-coordinating D133 state and a deprotonated, Cu-coordinating state (∼ 3.3-3.5 Å, Supplementary Fig. 3h), providing a possible structural basis for coupling proton transfer to the local Cu_C_ redox chemistry.

Together, these data support a bipartite catalytic model in which Cu_D_ is proposed to serve as the primary site for O_2_ activation and substrate hydroxylation, whereas Cu_C_ may mainly facilitate local methyl-PQ/methyl-PQH_2_ redox cycling. Such functional partitioning provides a plausible framework for coupling spatially separated oxygen activation and reductant regeneration within *Ni*AMO. This model differs from a recently proposed concerted Cu_C_–Cu_D_ catalytic model for AOB AMO ^18^ and raises the possibility of lineage-specific modes of copper-site utilization and quinone coupling.

### Structure of *Ni*AMO-inhibitor complex

To assess whether structural differences among AMOs translate into differential inhibitor susceptibility, we measured the IC_50_ values of five representative NIs against pure cultures of *N. inopinata* (Comammox), *N. maritimus* SCM1 (AOA) and *N. halophila* ZJB1 (AOB) (Fig. 3a, b and Supplementary Fig. 17).

**Fig. 3.**
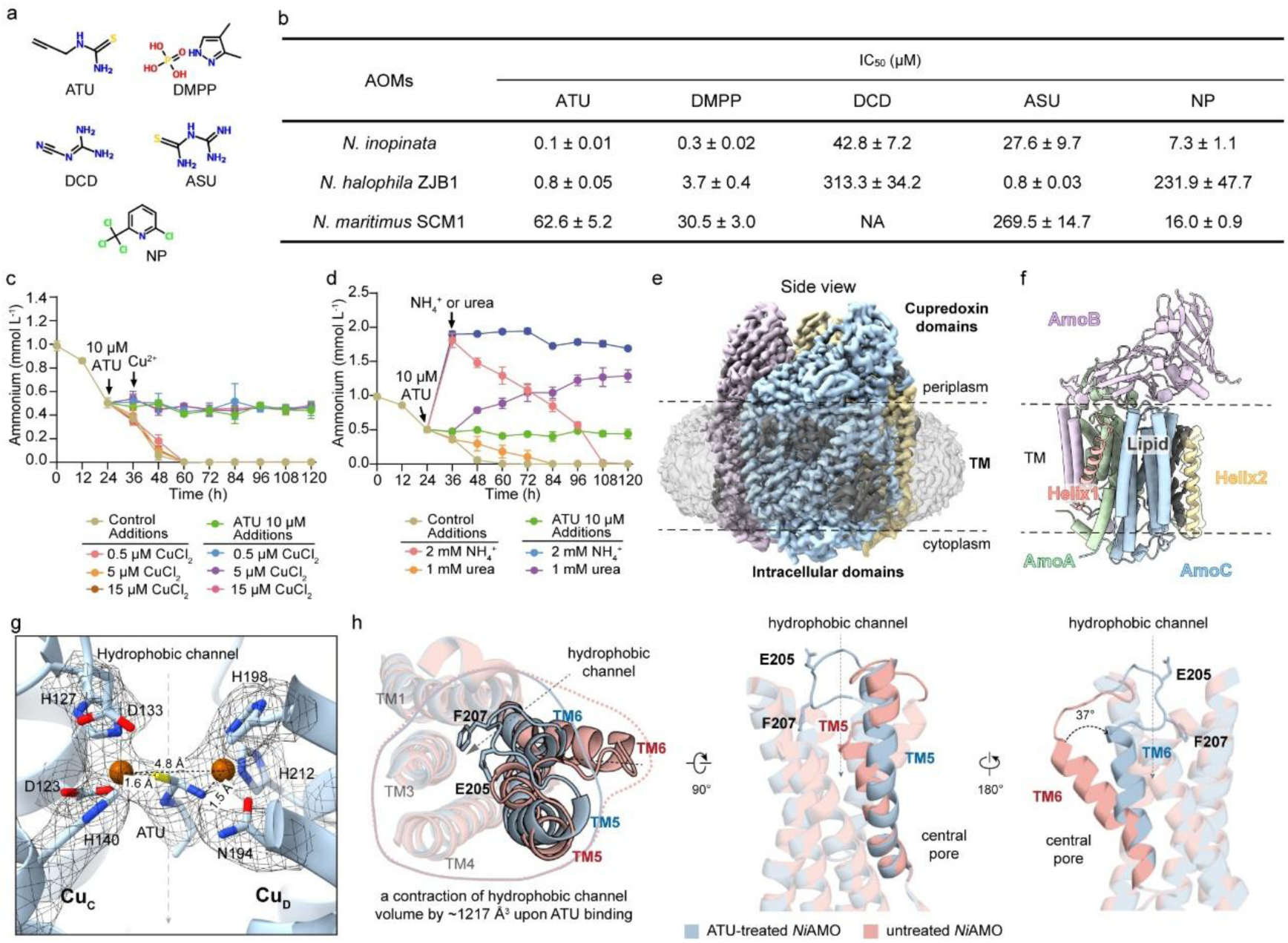
Overall structure of *Ni*AMO in complex with ATU. **a,** Chemical structures of five tested NIs. **b,** Differential inhibitory profile of NIs across phylogenetically distinct AOMs. IC_50_ values are expressed in micromolar (μM) and data are represented as mean ± SD (n = 3). NA: no inhibition detected at the tested concentration. **c-d,** Stress relief experiments on ATU-treated *N. inopinata* cells. Additions of copper **(c)** and ammonium and urea **(d)** to cultures treated with 10 μM ATU. Error bars represent standard deviations of the mean (triplicates). Black arrows indicate the time of addition of ATU, copper, ammonium, or urea to ATU-treated and control cultures. **e,** Cryo-EM density map of ATU-treated *Ni*AMO shown in side view. **f,** Atomic model of ATU-treated *Ni*AMO protomer, including AmoA (green), AmoB (light purple) and AmoC (blue), together with Helix1 (coral) and the newly resolved Helix2 (yellow). Lipid densities bridging Helix2 and the AMO core are depicted in gray. **g,** The simultaneous presence of Cu_C_ and Cu_D_ sites within the hydrophobic channel of inhibitor-bound *Ni*AMO. A well-defined ATU-derived density is positioned between Cu_C_ and Cu_D_. **h,** Structural comparison of hydrophobic channels between ATU-treated (blue) and untreated *Ni*AMO (coral).

The three phylogenetically distinct lineages displayed divergent susceptibility profiles, with IC_50_ values spanning more than three orders of magnitude. *N. inopinata* was the most sensitive of the three isolates to ATU, DMPP, DCD and NP at sub- to low-micromolar concentrations. *N. halophila* ZJB1 was highly sensitive to two thiourea derivatives ATU and ASU but comparatively resistant to DCD and NP. By contrast, *N. maritimus* SCM1 showed relatively low susceptibility to the inhibitors evaluated here (Fig. 3b and Supplementary Fig. 17), consistent with the previous observations^29^. Thus, *N. inopinata* showed the greatest inhibitor susceptibility among the three tested representatives. Its comparatively wider hydrophobic-channel opening, as revealed by structural analysis above, provides a mechanistic hypothesis for enhanced inhibitor access to the *Ni*AMO catalytic region.

Among the tested NIs, ATU showed the strongest inhibitory activity against *N. inopinata* (Fig. 3b). To elucidate the mechanistic basis of ATU-mediated inhibition, whole-cell assays were conducted (Fig. 3c, d). Exogenous CuCl_2_ failed to restore ammonia oxidation inhibited by ATU (Fig. 3c), indicating that ATU targets the intracellular copper center of AMO rather than scavenging bioavailable copper ions. Under ATU treatment, urea hydrolysis remained active and urea-derived ammonia was rapidly consumed in untreated controls (Fig. 3d), showing that ATU does not directly impair urease activity. Urea hydrolysis can therefore increase ammonia supply during ATU exposure, but it does not bypass the requirement for *Ni*AMO in subsequent ammonia oxidation. Partial relief of inhibition was observed upon supplementation with either NH_4_^+^ or urea-derived ammonia (Fig. 3d), consistent with substrate competition. These data support a copper-chelating, substrate-competitive mode of ATU inhibition in *N. inopinata* that differs from the recently observed in AOA^40^. To further resolve the structural basis of ATU inhibition, *Ni*AMO was purified from ATU-treated *N. inopinata* cells, and its cryo-EM structure was determined at 2.68 Å resolution (Fig. 3e and Supplementary Fig. 18). The overall structure of the *Ni*AMO-ATU complex closely resembles the inhibitor-free form (Fig. 3e, f and Supplementary Fig. 19a, b). A weak but discernible TM helix density adjacent to AmoC, hereafter Helix2, becomes visible in the inhibited state (Fig. 3f, Supplementary Figs. 4 and 19b). Helix2 comprises approximately 26 residues, lacks detectable periplasmic or cytoplasmic extensions, and packs against TM3, TM4 and TM5 of AmoC through lipid-mediated contacts (Fig. 3f). Its presence maybe reflect reduced conformational dynamics following ATU-induced inactivation.

In *Ni*AMO-ATU complex, no detectable ATU density was found in the Cu_B_ site (Supplementary Fig. 19c), supporting previous evidence that Cu_B_ site is not the catalytic center of AMO^15,17^. In contrast, a well-defined ATU-derived density was clearly resolved within the hydrophobic channel. ATU simultaneously chelated both Cu_C_ and Cu_D_ through its sulfur and nitrogen atoms, thereby stabilizing the Cu_D_ site into a defined conformation coordinated by N194, H198 and H212 (Fig. 3g and Supplementary Fig. 19d, e). ATU binding also enhanced the structural order of the TM5-TM6 loop. In this loop, E205 protrudes into the upper opening of the hydrophobic channel, where it forms a lid-like structure with F207, suggesting its potential role in regulating substrate access to the catalytic center (Fig. 3g, h and Supplementary Fig. 19e). Two newly formed hydrogen bonds from H212-F207 and N208-D51, further stabilize the conformation of this loop (Supplementary Fig. 19e).

Simultaneous chelation of Cu_C_ and Cu_D_ by ATU reduces the intermetallic distance to 4.8 Å and induces the conformational change in the surrounding helices that line the hydrophobic channel (Fig. 3g, h). Compared with the active *Ni*AMO structure, the upper segment of TM5 undergoes a subtle outward shift, whereas the upper segment of TM6 dramatically rotates inward by ∼37°, collectively reducing the hydrophobic channel volume (Fig. 3h). In addition, rearrangement of the Cu_C_–Cu_D_ center abolishes quinone-like density (Fig. 1e), indicating that inhibitor binding and hydrophobic channel closure to block access of the endogenous reductant to the catalytic center.

### Mechanism of ATU-mediated *Ni*AMO inhibition

Structural comparison of inhibitor-free and ATU-bound *Ni*AMO revealed that the conformational changes are concentrated around the Cu_C_–Cu_D_ region and the upper hydrophobic channel. In the inhibitor-free structure, the channel adopts an open configuration and contains a quinone-like density near the copper sites. In the ATU-bound structure, the inhibitor occupies the catalytic region, chelates both copper ions, contracts the hydrophobic channel and blocks reductant accessibility (Figs. 1 and 3). US simulations further showed that ATU could spontaneously access the *Ni*AMO, traverse a substrate channel with a minimum diameter of 6.7 Å en route to the Cu_C_ site (overcoming an energy barrier of 9.0 kcal/mol, Fig. 4a-c and Supplementary Fig. 20), and trigger structural reorganization of the Cu_D_ coordination environment into an ordered Cu_D_-bound configuration (Fig. 4d, e and Supplementary Fig. 21). This transition proceeds with a low activation barrier of 6.7 kcal/mol and is thermodynamically favorable (−4.2 kcal/mol), indicating that ATU binding both kinetically facilitates and thermodynamically stabilizes Cu_D_ coordination. These computational results are consistent with the well-resolved Cu_D_ density observed in the ATU-bound *Ni*AMO structure (Fig. 3g and Supplementary Fig. 19e).

**Fig. 4.**
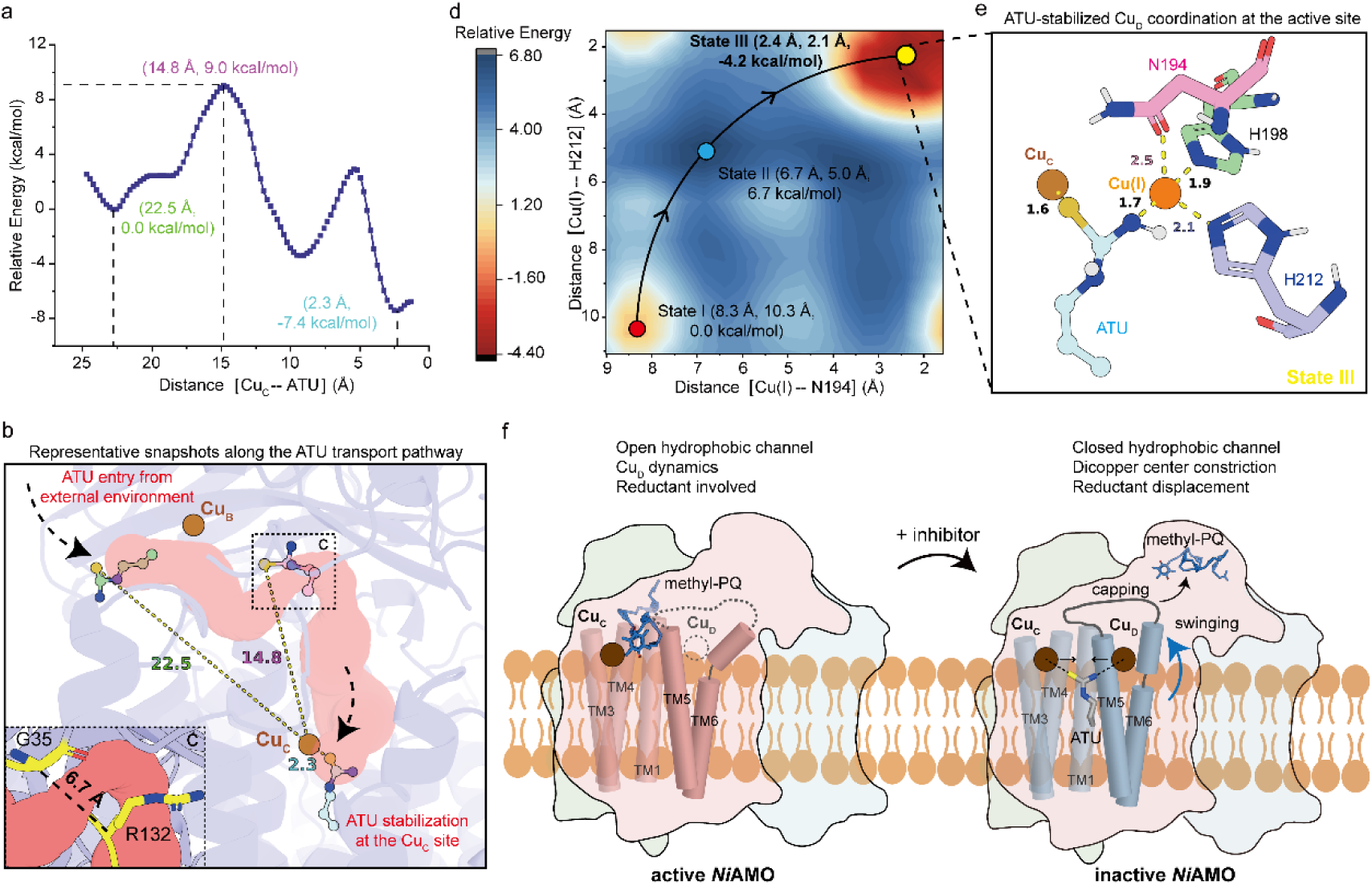
ATU-mediated inhibition mechanism on comammox *Ni*AMO. **a,** US-derived free-energy profile (kcal mol⁻^1^) for ATU migration from the extracellular environment into the active site of *Ni*AMO. The reaction coordinate was defined as the distance between the Cu(I) ion at the Cu_C_ site and the sulfur atom of ATU. **b-c,** Representative conformations of ATU along its migration pathway toward the active site. The representative ATU conformations are displayed as green, pink, and cyan ball-and-stick models, respectively (**b**). The narrowest region of the channel for ATU entry is enlarged in (**c**). The amino acid residues at the narrowest regions are displayed as yellow stick models. **d,** US-derived free-energy landscape (kcal mol⁻^1^) for the transition of Cu(I) from the initial non-coordinating configuration (State I) to the Cu_D_-coordinating configuration (State III). The free-energy landscape was defined by the distances of Cu(I) from N194 and H212. A representative structure illustrating the Cu_D_-coordinating state and the local coordination geometry of Cu(I) with N194 and H212 is shown in (**e**). Key interatomic distances are given in Å. **f,** Schematic illustration of ATU-mediated inhibition in comammox *Ni*AMO. In the active state, *Ni*AMO remains an open hydrophobic channel and a quinone-like density is positioned near this Cu_C_-Cu_D_ region. Upon ATU binding, the inhibitor bridges the Cu_C_ and Cu_D_ sites, induces rearrangement of the surrounding helices to restrict substrate access. Concurrently, the quinone-like density is displaced from the catalytic center. Copper ions are shown as spheres, and flexible or unresolved regions are indicated by dashed lines.

These observations support a coupled mechanism for ATU-mediated *Ni*AMO inhibition (Fig. 4f). First, ATU directly occupies the copper-containing cavity, preventing the catalytic region from adopting the dynamic configurations proposed for oxygen activation and substrate oxidation. Second, ATU-coupled rearrangements of TM5, TM6 and the TM5–TM6 loop promote formation of an E205/F207-containing lid and constrict the channel entrance, further restricting substrate access to the catalytic center. Third, the ATU-induced structural reorganization coincides with the loss of the quinone-like density, disrupting the local environment proposed to support electron and proton delivery.

ATU therefore inactivates *Ni*AMO through coordinated active-site occupation, channel closure and interference with electron and proton transfer. This multi-layered mechanism explains the high potency of ATU against *N. inopinata* and highlights the Cu_C_–Cu_D_ region, TM5-TM6 gating module and associated reductant-binding environment as interdependent determinants of *Ni*AMO inhibition.

### Multi-omics analysis link *Ni*AMO inhibition to an energy-limited cellular response

To probe cellular responses following direct ATU-mediated *Ni*AMO inhibition, we performed paired transcriptomic-proteomic profiling comparing untreated and ATU-treated *N. inopinata* after 12 h of exposure (Fig. 5). ATU treatment altered both transcript and protein pools, yielding 266 up- and 397 down-regulated transcripts, alongside 177 up- and 59 down-regulated proteins (Fig. 5a). Functional enrichment showed reduced representation of translation, ribosomal biogenesis and energy-production pathways, consistent with a broad downshift in biosynthetic and energy-conservation activity. By contrast, stress-sensing and signal-transduction increased at both transcript and protein levels (Fig. 5b and Supplementary Fig. 22), indicating a non-uniform cellular response rather than a generalized loss of function. Targeted inspection of nitrification-related genes revealed elevated *amoC* transcript abundance under ATU treatment. Together with the direct structural evidence for ATU binding, the absence of coordinated *amo* repression is consistent with inhibition of pre-existing *Ni*AMO rather than transcriptional shutdown of the enzyme complex (Fig. 5c). Several genes encoding respiratory complex I, ATP synthase and NirK were down-regulated, consistent with a broader decrease in energy-conservation and nitrogen-metabolism functions following *Ni*AMO blockade (Fig. 5c). Notably, urease and urea-transport components were up-regulated at both transcriptional and proteomic levels (Fig. 5c). As urea hydrolysis supplies ammonia but does not bypass AMO, we interpret this response as increased substrate-supply capacity compensate for competition under ATU exposure (Fig. 3d). Transcript-protein discordance for cell-division components, including accumulation of FtsB despite repression of cell-cycle transcripts, may reflect different response and turnover timescales and is not taken as evidence for preservation of an active divisome (Fig. 5c).

**Fig. 5.**
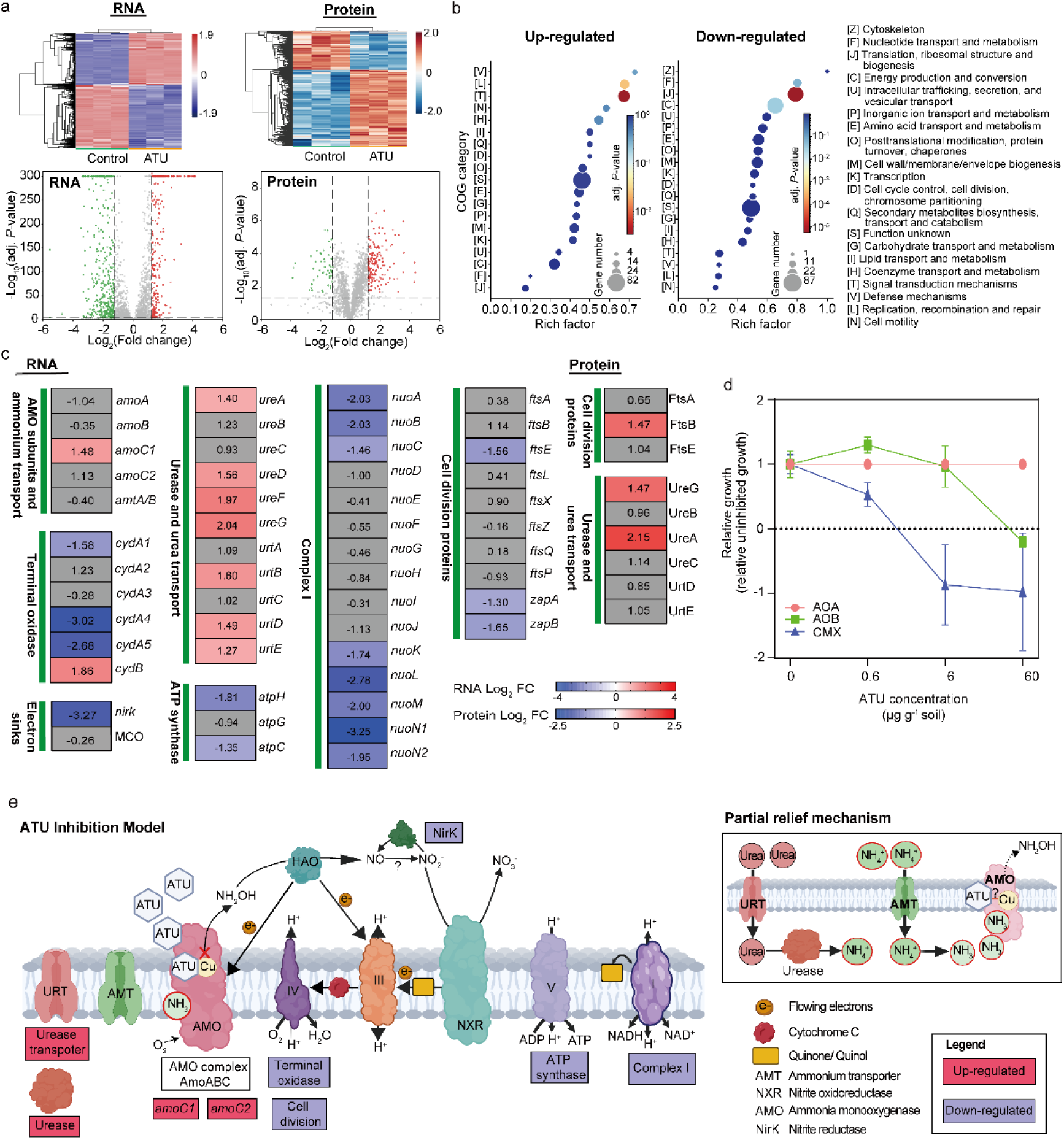
Multi-omics insights into ATU-mediated inhibition. **a,** Transcriptomic and proteomic heatmap of genes and proteins identified as significantly differentially expressed. Volcano plots of differentially expressed genes (DEGs) and proteins (DEPs) in response to ATU treatment. The x-axis shows log_2_(fold change) and the y-axis shows-log_10_(adjusted *P*-value). Thresholds: |log_2_FC| > 1.25 and adjusted *P* < 0.001 for DEGs;|log_2_FC| > 1.2 and adjusted *P* < 0.05 for DEPs. Full genes and proteins significantly changed can be found in Supplementary Data 1. **b,** Enrichment analysis of differentially expressed genes based on EggNOG database in every cluster from transcriptomic heatmap. **c**, Selected differentially expressed genes and regulated proteins. Full gene and protein annotations can be found in Supplementary Data 2. **d,** Response of soil ammonia oxidizers to ATU treatment. The y-axis shows a relative growth index calculated from *amoA* gene abundance in ATU-treated microcosms relative to untreated controls. A value ≤0 indicates complete inhibition, a value ≥1 indicates no inhibition or potential promotion of growth, and a value between 0 and 1 indicates partial inhibition. n=3 for biologically independent experiments; error bars indicate mean ± SD. **e**, Schematic model of cellular adaptive responses to ATU inhibition. ATU binds and chelates the copper-containing active site of *Ni*AMO, thereby directly blocking the initial ammonia-oxidation reaction and interrupting electron supply for downstream nitrification and respiration. The inset illustrates urea/ammonia-dependent stress-relief strategy. Red boxes mark up-regulated genes/proteins, and purple boxes denote down-regulated components. Key nitrogen-cycling enzymes, transporters and the electron-transport-chain complexes are depicted.

To assess whether differential ATU susceptibility extends to complex environmental communities, we quantified changes in *amoA* gene abundances of AOA, AOB and comammox *Nitrospira* in soil microcosms amended with graded ATU concentrations (Fig. 5d). AOA showed little suppression under the tested conditions, whereas comammox *Nitrospira* displayed greater suppression of net *amoA* gene accumulation than AOB (Fig. 5d), extending the differential ATU response observed in pure culture to this soil microcosm. Collectively, the structural, physiological and multi-omics data link direct *Ni*AMO inhibition to an energy-limited cellular response characterized by reduced biosynthetic functions, stress responses and increased urea-utilization capacity (Fig. 5e).

## Discussion

Complete nitrification integrates ammonia oxidation and nitrite oxidation within a single cell, yet the structural basis of the catalytic step that initiates this metabolism has remained unresolved. Here, high-resolution structures of inhibitor-free and ATU-bound *Ni*AMO, together with inhibition assays, MD and QM/MM simulations, and multi-omics data support a catalytic and inhibitory architecture that differs from that proposed for canonical AOB AMO. *Ni*AMO combines a distinct transmembrane architecture, TM6-dominated channel gating and a dynamic, methyl-PQH_2_-coupled Cu_D_ region with pronounced inhibitor susceptibility in *N. inopinata*.

*Ni*AMO contains two auxiliary transmembrane densities that occupy positions distinct from the accessory helices in reported AMO structures from AOB^16,17^. The current cryo-EM maps do not permit unambiguous sequence assignment, but mass spectrometry (Supplementary Fig. 3e) and genomic context^3,41^ suggest that these densities may derive from AmoD- or AmoE-like components. Helix1 could alternatively originate from a transmembrane segment associated with putative nitrite oxidoreductase (NXR), which catalyzes the second step of complete nitrification (Supplementary Fig. 3d-e)^3,42^. This possibility raises the testable hypothesis that AMO and NXR are spatially organized within a shared membrane microenvironment, but direct evidence for an AMO-NXR supercomplex, inter-enzyme electron transfer or obligatory substrate channeling will be required.

The most consequential divergence occurs within the Cu_C_–Cu_D_ region. High-resolution AOB AMO structures show the simultaneous occupancy of both Cu_C_ and Cu_D_ sites, and have been interpreted in terms of cooperative dicopper mechanism with markedly higher catalytic efficiency than either mononuclear Cu_C_ or Cu_D_ alone ^15,17,18^. In *Ni*AMO, although both Cu_C_ and Cu_D_ are implicated, O_2_ activation and substrate oxidation appear to occur predominantly at Cu_D_, whereas Cu_C_ mainly supports local quinone-quinol redox cycling rather than directly participating in the oxygenation chemistry. This functional partitioning supports a Cu_D_-centric catalytic pathway that differs from the concerted dicopper mechanism proposed for AOB AMO ^18^. Mechanistically, the two systems appear to diverge after formation of a Cu_D_–OOH^•⁻^ intermediate. In AOB AMO, the relatively close Cu_C_–Cu_D_ geometry permits Cu_D_–OOH^•⁻^ to engage Cu_C_ and evolve toward a bridging dicopper oxygen species, whereas the larger Cu_C_–Cu_D_ separation and steric constraints in *Ni*AMO favour further HAT and H_2_O_2_ formation at the mononuclear Cu_D_ site. Consistently, methyl-PQH_2_-mediated formation of the Cu_D_(II)– O^•⁻^ species in *Ni*AMO involves a rate-limiting H_2_O_2_ activation barrier of 17.5 kcal mol⁻^1^ (Supplementary Fig. 12), higher than the 14.7 kcal mol⁻^1^ barrier for the first HAT step that generates the Cu_D_(II)–OOH^⁻^ intermediate (rate-limiting step) in the CoQ_10_H_2_-coupled AOB AMO pathway ^18^. Taken together, the distinct fate of the Cu_D_(II)–OOH^⁻^ intermediate and the resulting higher-barrier mononuclear oxygen-activation pathway may contribute to the substantially lower substrate turnover rate observed in comammox bacteria relative to canonical AOB.

The widened, TM6-regulated channel entrance provides a structural rationale for the high ATU sensitivity of *N. inopinata* relative to the AOB isolate tested here. US indicates that ATU can traverse the channel to the Cu_C_ region over a free-energy barrier of approximately 9.0 kcal mol^-1^. Once bound, ATU does more than compete for space: it bridges Cu_C_ and Cu_D_, traps the otherwise dynamic Cu_D_ coordination environment, orders the E205/F207-containing lid, constricts the channel and displaces the quinone-like density. Thus, the high sensitivity may arise from the combination of accessible inhibitor entry and conformational trapping rather than copper chelation alone. Given the broad distribution of comammox^12^, this mechanistic divergence is ecologically relevant. Sequence analysis shows that key structural elements are highly conserved in comammox clades A and B, including the TM5–TM6 region and Cu_C_–Cu_D_ coordination environment (Fig. 1g), suggesting these structural features may be conserved across the comammox lineage. A broader generalization to comammox requires additional isolates and environmental populations.

The whole-cell rescue assays complement the structural observations by supporting substrate competition between ammonia and ATU for access to the *Ni*AMO catalytic region. Exogenous copper supplementation failed to reverse ATU-mediated ammonia-oxidation suppression, arguing against extracellular copper sequestration and supporting direct inhibition of the intrinsic copper-containing site. Partial relief by excess ammonium or urea-derived ammonia is also consistent with competition for access to *Ni*AMO, a response distinct from that recently reported in AOA^40^. Consistent with this physiological phenotype, multi-omics profiling showed that ATU treatment reduces translation, ribosomal-biogenesis and energy-production functions, consistent with an energy-limited maintenance response, whereas increased stress-signal functions indicate active physiological reprogramming. The concordant increase in urea-transport and urease components is particularly informative: urea hydrolysis cannot bypass *Ni*AMO, but it can increase ammonia supply and thereby provide a plausible compensatory response under competitive ATU inhibition. The soil microcosm data further indicated that, under the tested conditions, comammox *Nitrospira* gene accumulation is more strongly suppressed than that of AOB. These observations connect the atomic-level inhibitory architecture to cellular and community-level responses, and also imply that urea-driven ammonia supply in soil environments may mitigate ATU inhibition of comammox populations under field conditions.

The *Ni*AMO structure also has practical implications for nitrogen management. AMO controls the entry of fertilizer-derived ammonium into nitrification and thereby influences nitrate formation, nitrogen retention and downstream N_2_O production. Comammox contribute to nitrification in agricultural soils^32,34^, and *N. inopinata* can grow on guanidine, with comammox-containing soil microcosms indicating that guanidine-derived nitrogen can enter environmental nitrification^14^. Inhibitors developed primarily against AOB may therefore not predictably suppress comammox. The widened *Ni*AMO channel, the Cu_C_–Cu_D_ region and the quinone-access pathway identify three testable determinants for lineage-aware inhibitor design or selection: channel access, copper coordination and disruption of electron/proton delivery. Translation to field application will require validation in soils with quantified comammox activity, together with measurements of nitrogen retention, N_2_O emissions and effects on non-target nitrifiers.

Together, these findings show that *Ni*AMO uses a distinct methyl-PQH_2_–Cu_D_ catalytic module and an ATU-responsive gating mechanism, providing a molecular basis for interpreting lineage-specific nitrification in ecosystems and for developing inhibitor strategies that explicitly account for comammox.

## Methods

### Cultivation of ammonia-oxidizing microorganisms

Three ammonia-oxidizing strains—*Nitrospira inopinata*, *Nitrosopumilus maritimus* SCM1, and *Nitrosomonas halophila* ZJB1—were cultured as previously described^3,4,17^. Briefly, pure cultures of *N. inopinata* were grown in modified AOM medium containing (per liter) KH_2_PO_4_ (0.05 g), KCl (0.075 g), MgSO_4_·7H_2_O (0.05 g), NaCl (0.58 g), CaCO_3_ (4 g), together with 1 mL non-chelated trace element solution (TES) and 1 mL selenium-wolfram solution (SWS)^3^. Following autoclaving, the initial pH reached approximately 8.2, and was subsequently maintained near 7.8 during pure-culture growth via the CaCO_3_-based buffering system^3^. *N. maritimus* SCM1 was cultivated in modified HEPES-buffered synthetic Crenarchaeota medium containing (per liter) NaCl (26 g), MgSO_4_·7H_2_O (5 g), MgCl_2_·6H_2_O (5 g), CaCl_2_·2H_2_O (1.5 g), KBr (0.1 g), 10 mL HEPES buffer (1 mol L^-1^ HEPES with 0.6 mol L^-1^ NaOH, pH 7.5), 5 mL KH_2_PO_4_ solution (0.4 g L^-1^), 2 mL bicarbonate solution (1 mol L^-1^), 1 mL FeNaEDTA solution (7.5 mol L^-1^), and a mixture containing 1 mL vitamin solution, 1 mL TES, streptomycin (100 mg mL^-1^), and amphotericin B (1 mg mL^-1^)^4,41,43^. *N. halophila* ZJB1 was maintained in a modified synthetic seawater medium containing (per liter) NaCl (5 g), KCl (0.25 g), MgSO_4_·7H_2_O (0.04 g), CaCl_2_·2H_2_O (0.1 g), 5 mL KH_2_PO_4_, 1 mL TES and 1 mL phenol red (0.05%) serving as pH indicator. The pH was adjusted to 8.0 using 10% Na_2_CO_3_^17,44^. Stock cultures were grown in 2 L bottles containing 1.2 L of medium and were supplied with NH_4_^+^ as the nitrogen source at 3 mmol L^-1^, 3 mmol L^-1^ and 10 mmol L^-1^ for *N. inopinata*, *N. maritimus* SCM1, and *N. halophila* ZJB1, respectively. *N. inopinata* and *N. maritimus* SCM1 were incubated without agitation at 42 °C and 30 °C, respectively, whereas *N. halophila* ZJB1 was grown with shaking at 120 rpm at 30 °C. To prevent photoinhibition, all strains were kept in darkness. Culture progression was monitored by measuring NH_4_^+^ consumption and culture purity culture purity was assessed by inoculation into LB broth according to a previously described method^45^.

### Inhibitor activity assays

The inhibitory responses of AOMs to five commonly used fertilizer additives— allylthiourea (ATU), 3,4-dimethylpyrazole phosphate (DMPP), dicyandiamide (DCD), amidinothiourea (ASU), and nitrapyrin (NP)—were investigated to evaluate the efficacy of NIs against AOMs. For each assay, exponentially growing AOM cells were collected from batch cultures and dispensed into sterile 100 mL glass bottles at 20 mL per bottle, after which the medium was supplemented with the inhibitor of interest across a graded concentration series. Control treatments consisted of identically inoculated cultures lacking NIs. Samples were collected at regular intervals 6 or 12 h to monitor NH_4_⁺ consumption, thereby tracking AO activity in AOMs.

### Chemical analyses

Ammonium was determined using the optimized indophenol blue method^46^. In brief, 200 μL aliquots of culture were transferred into sterile 1.5 mL microcentrifuge tubes and centrifuged at 12,000 rpm for 5 min at room temperature to pellet cellular debris and large particles. The resulting supernatant was transferred to fresh sterile 2 mL Eppendorf tubes and diluted to a final volume of 1 mL with NH_4_^+^-free ultrapure water. For the optimized indophenol blue method, 50 μL of phenol-nitroprusside solution and 50 μL of alkaline hypochlorite solution were added. Once color development had reached completion, absorbance was recorded at 630 nm on an EnSpire microplate reader. Final concentrations were calculated using standard curves for NH_4_^+^ (0-10 μM, *n* = 3, R^2^ > 0.998).

### Membrane fraction extraction and AMO protein purification

For membrane preparation, *N. inopinata* cells were cultured in modified AOM medium either without additive (for active AMO) or in the presence of 10 μM ATU (for inactivated AMO). 80 L of *N. inopinata* cells were harvested by vacuum filtration through a 0.22 µm pore polycarbonate membrane filter (Millipore) and resuspended in 50 mL lysis buffer (50 mM PIPES pH 7.2, 50 mM NaCl, 30 μM CuSO_4_); for inactivated AMO preparation, the lysis buffer was supplemented with 10 μM ATU throughout. Cell disruption was achieved via ultrasonication (Miou Instrument, Zhejiang, China) at 80% output using a 2 s-on/4 s-off duty cycle over a total of 30 min. The resulting lysate was centrifuged at 12,000 rpm at 4 °C for 30 min to remove cellular debris and intact cells. The supernatant was collected and ultracentrifuged at 150,000 ×g at 4 °C for 2 h. The extracted membrane pellet was resuspended in 100 μL of lysis buffer, then centrifuged at 13,000 rpm and 4 °C for 20 min. The supernatant was discarded, and 100 μL of lysis buffer was added to resuspend the membrane pellet. This washing procedure was repeated three times. Finally, the membrane pellet was resuspended in 40 μL of lysis buffer. Subsequently, n-Decyl-β-D-maltopyranoside (DDM) was added to a final concentration of 0.5%, and the mixture was stirred at 4 °C for 2 h to extract membrane proteins. Afterward, the sample was centrifuged at 13,000 rpm and 4 °C for 10 min to remove cell debris. The supernatant was collected and subjected to ultracentrifugation at 150,000×g for 2 h at 4 °C to remove ribosomes. The supernatant containing AMO proteins was collected, flash-frozen in liquid nitrogen, and stored at -80 °C until further use.

### Preparation of cryo-EM samples and data acquisition

Purified samples were first evaluated by negative-stain electron microscopy on a Talos L120C G2 instrument (Thermo Fisher Scientific) operated at 120 kV. For vitrification, holey-carbon gold grids (Quantifoil R1.2/1.3, 300 mesh) overlaid with a monolayer of graphene (a gift from Jiayue Su, Tsinghua University) were subjected to glow discharge, after which 4 µL of samples was applied onto the front side of the plasma-cleaned grids. Vitrification was accomplished by plunge-freezing into liquid ethane using a Mark IV Vitrobot (ThermoFisher) maintained at 4 °C under 100% relative humidity. Cryo-EM grids were subsequently transferred into liquid nitrogen for storage until data collection. The vitrified samples were loaded into a Titan Krios electron microscope G4 (Thermo Fisher Scientific) operated at 300 kV, equipped with Selectris energy filter (10eV slit width) together with a CEOS CCOR-spherical aberration/chromatic aberration (Cs/Cc) corrector. Automated data collection yielded 15,188 and 8,319 movie stacks for the active and ATU-bound NiAMO datasets, respectively, all recorded on a Falcon 4 detector operated in counting mode at a nominal magnification of ×130,000 (calibrated pixel size of 0.6584 Å). Each movie was dose-fractionated into 40 frames, with a total dose of 50 ^e-^/Å^2^. Data acquisition was performed using EPU2 software package with a defocus range from -1.2 to -2.0 μm. More data collection details are given in Supplementary Table 4.

### Image processing

All raw micrographs were first corrected for beam-induced motion using MotionCor2^47^. The cryoSPARC v4.3.2^48^ was used for all downstream image processing. Contrast transfer function (CTF) determination was performed via the Patch CTF routine, and suboptimal micrographs identified by manual inspection were excluded. For particle picking, the Blob Picker was applied using an 8-to-20-nm diameter search window; picked particles were boxed at 224 pixels and Fourier-cropped by a factor of 2. Two-dimensional (2D) classification was then carried out iteratively, and high-quality 2D averages were fed into Topaz^49^ for automated particle picking. This two-step procedure—classification followed by Topaz training and re-picking—was cycled four times until 2D class averages ceased to improve. All particles collected across these iterations were pooled, and duplicated entries were eliminated. The resulting non-redundant particle set was used for ab initio volume generation, followed by heterogeneous refinement, which produced several class volumes. The volume exhibiting features of the AMO complex was subsequently refined against prior maps to remove spurious or low-occupancy particles. For the active NiAMO dataset, after three rounds of this reference-based sorting, the best class was re-processed through a fresh ab initio job, then two more heterogeneous refinement steps, and finally subjected to NU refinement with C3 symmetry imposed^50^, yielding a 2.47-Å map. For the ATU-treated *Ni*AMO sample, following the same three sorting rounds, the data were directly advanced to NU refinement (C3)^50^, which produced a 2.68-Å reconstruction. DeepEMhancer^51^ was finally applied to all refined maps to improve density quality for modeling. As a control, the full processing pipeline was also executed without symmetry (C1), which resulted in nearly identical maps with only a marginal loss of resolution.

### Model building and refinement

Initial atomic models for NiAmoA, NiAmoB, and NiAmoC were generated de novo using AlphaFold3^52^ and rigid-body fitted into the cryo-EM density maps using UCSF Chimera^53^. Subsequent manual model rebuilding was performed in Coot^54^, followed by iterative real-space refinement in PHENIX^55^ with secondary structure and geometric restraints enforced. The two auxiliary transmembrane helices were modeled as polyalanine due to insufficient side-chain density. Model geometry and map-to-model agreement were evaluated using standard cryo-EM validation metrics. Comprehensive model-building and validation statistics are provided in Supplementary Table 4.

### Mass spectrometry analysis

To further characterize the composition of the CMX-AMO complex, the purified AMO samples were resolved on a 4-20% SDS-PAGE gel (Yeasen Biotechnology). The gel bands corresponding to the AMO fractions were then excised and transferred into sterile 1.5 mL microcentrifuge tubes. These gel pieces were subsequently washed four times with ultrapure water to remove residual SDS-PAGE running buffer. Finally, the washed AMO protein bands were subjected to LC-MS/MS analysis.

Liquid chromatography-tandem mass spectrometry (LC-MS/MS) analysis was performed on a Thermo-Dionex Ultimate 3000 HPLC system coupled to an Orbitrap Fusion mass spectrometer (Thermo Scientific). Peptide separation was achieved over a 120-min gradient at 0.300 µL min^-1^ on a homemade 150-mm C18 column (75 µm i.d.; 5 µm resin; 300 Å). The mobile phase comprised 0.1% formic acid (A) and 0.1% formic acid with 100% acetonitrile (B). Data acquisition was carried out in data-dependent mode via Xcalibur 3.0, with full MS scans collected in the Orbitrap (350-1,550 m/z; 120,000 resolution) followed by 3-s MS/MS scans at 30% normalized collision energy in the ion-routing multipole. The MS/MS spectra were searched against the designated database using Proteome Discoverer (v.1.4).

### Copper quantification by inductively coupled plasma mass spectrometry (ICP-MS)

Purified AMO samples were dried in a vacuum centrifugal concentrator (Jiaimu, Beijing, China) and accurately weighed (0.0001 g) portions were transferred into acid-cleaned polytetrafluoroethylene digestion vessels. Samples were pre-digested at room temperature for 30 min with nitric acid and hydrochloric acid (6 ml HNO_3_ and 2 ml HCl), then subjected to a stepwise hotplate digestion (100 °C/10 min to 120 °C/10 min to 200 °C/60 min), with acid replenished as needed to prevent drying. After complete digestion, the solution was gently evaporated to ∼0.5 mL to remove excess acid, cooled, and made up to 10 mL with metal-free water in a volumetric flask; the vessel was rinsed three times and rinses combined. Copper content was quantified by Agilent 7500CE ICP-MS instrument (Agilent Technologies, USA). The metal content was normalized to the AMO protomer concentration to calculate copper stoichiometry (Supplementary Table 5).

### Relief experiments

Following the identification of differentially expressed genes and proteins, 1 L of *N. inopinata* culture was cultivated and supplemented with 1 mM NH_4_^+^. After 24 h of incubation, the culture was aliquoted into 20 mL portions in 50 mL sterile plastic centrifuge tubes, and divided into ATU-treated (final concentration 10 μM) and control groups. Following 12 h of ATU exposure, the following reagents were respectively added to assess the recovery capacity from inhibitor-induced stress: CuCl_2_ (0.5, 5 and 15 μM), NH_4_Cl (2 mM), and urea (1 mM). Each condition was performed with three biological replicates. Ammonium concentrations were monitored at 6 h intervals to track the ammonia-oxidizing activity of *N. inopinata*.

### Transcriptomic and proteomic analyses

A total of 60 L cultures of *N. inopinata* were grown. Half were maintained as untreated controls, whereas the other half were treated with 10 μM ATU for 12 h. Following treatment, cells from each group were harvested by vacuum filtration onto 0.22-μm pore-size membranes (Millipore), with three biological replicates per condition. The resulting cell pellets were immediately flash-frozen in liquid nitrogen and stored at - 80 °C prior to downstream omics analyses.

For transcriptomic profiling, total RNA extracted from growing *N. inopinata* samples was sent to Magigene Biotechnology Co., Ltd. (Guangzhou, China) for cDNA library construction and high-throughput sequencing on an Illumina HiSeq 2500 platform. Differential gene expression between the CMX-control and CMX-ATU (10 μM ATU) groups was evaluated using the DESeq2 R package^56^ based on normalized read counts. Transcripts satisfying the criteria of |log_2_(fold change)| > 1.25 and adjusted *P-*value < 0.001 were classified as differentially expressed genes (DEGs). Functional annotation of the assembled unigenes was performed via BLAST searches against the NR database (https://www.ncbi.nlm.nih.gov/). In the R environment, functional enrichment analysis of DEGs was conducted based on the EggNOG database^57^.

For quantitative proteomic profiling, an 8-plex iTRAQ reagent kit (AB SCIEX, USA) was applied following a previously described protocol^58^ with minor modifications. The iTRAQ-labeled peptide mixtures were initially fractionated by high-pH reversed-phase HPLC on an Agilent 1260 series system fitted with an Agilent 300 Extend C18 column. Peptide separation was then accomplished on an EASY-nLC 1000 UPLC system (ThermoFisher Scientific) equipped with an Acclaim PepMap RSLC reversed-phase analytical column, and tandem mass spectrometry (MS/MS) was acquired on a Q Exactive Hybrid Quadrupole-Orbitrap mass spectrometer (ThermoFisher Scientific). Proteins meeting the thresholds of |log_2_(fold change)| > 1.20 and adjusted *P-*value < 0.05 were regarded as differentially expressed proteins (DEPs). Functional annotation and enrichment analyses were performed for the identified proteins. Full details of the transcriptomic and proteomic analyses are available in Supplementary data 1 and 2

### Microcosm incubation

To evaluate the sensitivity of ammonia-oxidizing microorganisms to ATU in agricultural soil, a 30-day soil microcosm incubation experiment was conducted. Surface soil (0-5 cm depth) was collected from M1 site (21.6442°N, 110.9139°E) and subjected to two treatment categories: 100 mg NH_4_^+^-N kg^-1^ soil + water (control), and 100 mg NH_4_^+^-N kg^-1^ soil + ATU (0.6, 6 and 60 μg g⁻¹ soil). Each treatment was established with three biological replicates. Fresh soil (∼20 g, moisture content ∼7.0%) was dispensed into 120 mL serum bottles, and solutions containing ammonium sulfate and ATU were thoroughly mixed with the soil to adjust the water-filled pore space (WFPS) to 50% of maximum water-holding capacity^59^. The bottles were then sealed with black butyl rubber stoppers and incubated at 25 °C in the dark. The stoppers were opened every 3 days for aeration, and the WFPS was maintained at its initial level by weekly addition of ultrapure water. Soil samples were collected on days 0 and 30, immediately frozen in liquid nitrogen and stored at -80°C until DNA extraction.

### DNA extraction and quantification of *amoA* gene

Soil DNA was extracted from ∼ 0.25 g of the soil using the Soil Genomic DNA Extraction Kit (Tiangen Biotech. Co., Ltd., Beijing, China). The DNA concentrations were measured using the NanoValue™ Plus micronucleic acid analyzer (GE Healthcare Life, Pittsburgh, PA, USA). The extracted DNA was stored at -20 °C for subsequent analyses.

The abundances of AOA, *β*-AOB and comammox clade A were quantified by quantitative PCR (qPCR) targeting archaeal and bacterial *amoA* genes, respectively, using the primers listed in Supplementary Table 6. qPCR reactions were performed on a QuantStudio 5 Real-Time PCR system (Applied Biosystems, CA, USA). Each 20-μL reaction contained 10 μL ChamQ™ Universal SYBR qPCR Master Mix (Vazyme, Nanjing, China), 0.5 μL forward primer, 0.5 μL reverse primer, 1 μL DNA template, and nuclease-free water. All samples and standard reactions were analyzed in triplicate

### Evaluation of ATU efficacy on soil ammonia-oxidizing communities

The inhibitory efficacy of ATU on specific ammonia-oxidizing communities in soil was evaluated by comparing community growth in inhibitor-amended microcosms relative to that in unamended controls. Relative growth rate was calculated using the following formula:

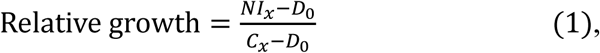

where “*NI_x_*” represents gene abundance on day x (day 30) in the presence of the inhibitor, “*C_x_*” represents gene abundance on day x in the control microcosms without any inhibitor, and “*D_0_*” represents gene abundance measured at day 0. If the value of relative growth is ≤ 0, it suggests complete inhibition by the ATU inhibitor; a value ≥ 1 indicates non-inhibition or promotion of growth by the ATU inhibitor; and a value between 0 and 1 indicates partial inhibition.

### System setup and molecular dynamics simulations

All molecular dynamics (MD) simulations were performed using the Amber Package, version 22 (University of California, San Francisco, CA)^60^. The initial models of *Ni*AMO determined in this study were employed. For the titratable residues (His, Asp, Glu), protonation state was assigned based on pKa values calculated in the PlayMolecule website (https://open.playmolecule.org)^61^ and detailed visual examination of the regional hydrogen-bonded networks. Within the *Ni*AMO monomer, H42 and H44 (in *Ni*AmoA), and H34, H137, H138 and H193 (in *Ni*AmoB) were protonated at the *δ* position, whereas H23, H172, H215 and H270 (in *Ni*AmoA), H86, H136, H140, H259, H288, H364 and H382 (in *Ni*AmoB), and H43, H127, H140, H198 and H212 (in *Ni*AmoC) were protonated at the *ε* position. For Asp and Glu residues, D25 (in *Ni*AmoA) was protonated, while the remaining residues were deprotonated. The membrane environment of *Ni*AMO was constructed using the Packmol-Memgen program^62^, with the phospholipid bilayer consisting of 60% phosphatidylvinylethanolamine (PVPE), 20% phosphatidylvinylglycerol (PVPG), and 20% 1-palmitoyl-2-oleoyl-sn-glycero-3-phosphocholine (POPC). The Amber ff14SB^63^, Lipid21^64^, and GAFF^65^ force fields were applied to parameterize the amino acid residues within *Ni*AMO, phospholipid bilayer, and reductants methyl-PQH_2_, respectively, while the force field parameters for the Cu_B_, Cu_C_ and Cu_D_ sites were tailored using the “MCPB.py” tool^66,67^ of AmberTools23. The membrane-embedded simulation system was solvated with a periodic rectangular box (the volume is 144.593 × 144.593 × 152.067 Å^3^) containing 68,853 TIP3P water molecules, 579 lipid molecules, and 0.15 M NaCl.

Following appropriate setup, the complex systems were successively fully minimized by combining 10,000 steps of the steepest-descent method and 10,000 steps of the conjugate gradient method. Then, each system was gradually heated from 0 K to 300 K for a total of 50 ps using the NVT ensemble. To achieve a uniform density after heating dynamics, 1 ns of density equilibrium was executed under the NPT ensemble, during which the temperature and pressure of the system were maintained at 300 K and 1.0 ATM using the Langevin thermostat and the Berendsen barostat^68,69^, respectively. Subsequently, all complexes were equilibrated for 2 ns without any restraints in order to relieve minor unfavorable ligand-protein steric interactions that might have been still present. Finally, a production MD simulation of 200 ns was carried out under the NPT ensemble. Throughout the process, the simulation integration step was set to 2 fs, and the conformational configurations of the complex systems were saved every 100 ps in trajectory files. The covalent bonds connecting hydrogen atoms were then restricted with the SHAKE algorithm^70^. The short-range nonbonded interactions were adopted using a cutoff radius of 8 Å, while the long-range electrostatic interactions were modeled utilizing the Particle mesh Ewald (PME) method with a grid point density of 0.1 nm and an interpolation order of 4^71^. The results of MD simulations were presented by using the Visual Molecular Dynamics version 1.9.3a^72^, and PyMOL software was used to exhibit the graphics of the MD simulations.

### Umbrella sampling

Umbrella sampling simulations were performed to characterize the free-energy landscapes associated with the Cu ion capture at the Cu_D_ site and ATU translocation into the active site of *Ni*AMO. Two-dimensional umbrella sampling simulations were conducted for mPQH_2_-free *Ni*AMO, mPQH_2_-bound *Ni*AMO, and ATU-bound *Ni*AMO structures to evaluate the thermodynamic feasibility of Cu ion recruitment and stabilization at the Cu_D_ site. The two-dimensional collective variables were defined as the distances between the Cu ion of Cu_D_ site and the coordinating residues N194 and H212, respectively. In addition, one-dimensional umbrella sampling simulations were carried out to determine the free-energy profile of ATU translocation from the external solvent environment into the active site of *Ni*AMO, with the reaction coordinate defined as the distance between ATU and the Cu ion at the Cu_C_ site. Umbrella sampling windows were generated along each reaction coordinate at 0.2 Å intervals. For each window, 10 ns of MD simulations were performed under a harmonic biasing potential with a force constant of 50 kcal/mol/Å^2^. The unbiased potentials of mean force (PMFs) were then reconstructed using the weighted histogram analysis method (WHAM)^73^, yielding quantitative free-energy landscapes for Cu ion capture at the Cu_D_ site and ATU translocation into the *Ni*AMO active site.

### QM/MM-MD and metadynamics simulations

All QM/MM Born−Oppenheimer MD simulations were performed utilizing the CP2K program version 2024.1 (https://www.cp2k.org). The initial structures for QM/MM MD simulation were derived from the representative conformations extracted from the classical MD trajectories. The interaction energies of the QM and MM regions were calculated separately using QUICKSTEP^74^ and the FIST module within the CP2K program, and the real-space multigrid technique^75^ was employed to calculate the electrostatic coupling between the QM and MM regions. The QM region was processed by using a mixed Gaussian and plane wave (GPW) basis set at the DFT (B3LYP) level^76^, and the MM region was parametrized in the same way as the classical MD simulation. For the conformational rearrangement of the methyl-PQH^•^ (mPQH^•^) radical near the Cu_D_ site, the QM region comprised the Cu_D_(II)–OOH^−^ species, the side chains of N194, H198, and H212, and the polar headgroup moiety of the mPQH^•^ radical. For the conformational rearrangement of mPQH^•^ near the Cu_C_ site, the QM region included the Cu(I) ion of the Cu_C_ site, the side chains of D123, H127, H140, protonated D133, and the aromatic ring moiety of the mPQH^•^ radical. All QM atoms at the boundary employed hydrogen-capped atoms to fill the empty valence. The wave function was expanded using a Gaussian double-*ζ* valence-polarized basis set (DZVP)^77^. An auxiliary plane-wave basis set with a cutoff energy of 360 Ry was implemented to converge the electron density, and combined with the Geodecker−Teter−Hutter (GTH) pseudopotential to handle the core electrons^78^. All QM/MM-MD simulations of 50 ps were performed under the NVT system using 0.5 fs integration steps and the simulated temperature was controlled by canonical sampling through velocity rescaling (CSVR) with a time constant of 10 fs^79^.

Finally, well-tempered metadynamics simulations^80,81^ were performed to characterize the free-energy profiles associated with the conformational rearrangement of the mPQH • radical near the Cu_C_ and Cu_D_ sites, respectively. For the mPQH^•^ radical conformational rearrangement near the Cu_D_ site, the collective variable (CV) was defined as the difference between the two distances connecting the oxygen atoms of the mPQH^•^ radical and the oxygen atom of the Cu_D_(II)–OOH⁻ specie. For the corresponding rearrangement near the Cu_C_ site, the CV was defined as the difference between the two distances connecting the oxygen atoms of the mPQH^•^ radical and the oxygen atom of protonated D133. Gaussian-shaped potential hills were defined by a width of 0.1 Å, and a height of 0.6 kcal/mol, with a deposition interval of 10 fs.

### QM/MM calculations

The last snapshot extracted from MD or QM/MM MD trajectories was used for the QM/MM calculations. All QM/MM calculations were performed using ChemShell^82,83^, combining turbomole^84^ for the QM region and DL_POLY^85^ for the MM region. The electrostatic embedding scheme^86^ was used to account for the polarizing effect of the protein environment on the QM region. Hydrogen link atoms with the charge-shift model were applied to treat the QM/MM boundary. During QM/MM geometry optimizations, the QM region was studied with the hybrid UB3LYP^87–89^ density functional with two levels of theory. For geometry optimization, the double-*ζ* basis set def2-SVP was used. The energies were further corrected with the larger def2-TZVP basis set for all atoms. Dispersion corrections computed with Grimme’s D3 method^90–92^ were included in all QM calculations. Candidate transition regions were identified from finely scanned potential-energy surfaces along the specified reaction coordinates using an increment of 0.02 Å, and all minima were optimized without symmetry restraints using the DL-FIND95 optimizer^93^. For the first stage of the H_2_O_2_ formation, QM/MM calculations were performed for both the open-shell singlet (OSS) and triplet spin states. The QM region was defined to include the polar headgroup moiety of methyl-PQH_2_ (mPQH_2_), the Cu_D_(II)–superoxo species, and the directly coordinating residues N194, H198, and H212. For the subsequent generation of the Cu_D_(II)–O^•−^ species, the reaction was investigated in the singlet state, with N190 additionally incorporated into the QM region. For the methyl-PQ (mPQ) reduction to mPQH_2_ near the CuC site, QM/MM calculations were performed to characterize the sequential proton-coupled electron transfer (PCET) processes. The QM region was defined to include the Cu(I) ion at the Cu_C_ site, the directly coordinating residues D123, H127, and H140, the protonated residue D133, and the polar headgroup moiety of mPQ. The first PCET step was optimized in the open-shell singlet state, while the second PCET step was investigated in the doublet state. The computed relative energies were electronic energies from the QM/MM calculation. The previous work in other metalloenzymes confirmed that the electronic energy barrier is close to the free energy barrier^94–99^. The spin density of QM/MM simulated species are shown in Supplementary Table 3.

### QM calculations

To evaluate the reaction mechanisms and energetics of the Cu_D_ site in the oxidation of NH_3_ to NH_2_OH (Supplementary Fig. 13), density functional theory (DFT) calculations were carried out using the Gaussian 16 software package (https://gaussian.com/citation/). Geometry optimizations were performed at the B3LYP/def2-SVP levels of theory in conjunction with the SMD implicit solvation model^72^, followed by single-point energy refinements at the B3LYP/def2-TZVP levels. Given that the *Ni*AMO active site is embedded within the protein–membrane environment, chlorobenzene was selected as the solvent in the SMD model^72^ to approximate the hydrophobic milieu. Dispersion corrections computed with Grimme’s D3 method were included in all QM calculations^91,92^, with all reported energies further refined using the larger def2-TZVP basis set for all atoms. The energy data are shown in Supplementary Table 2, and the spin density data are shown in Supplementary Table 3.

## Data availability

The cryo-EM maps of purified *Ni*AMO complex without (EMD-83338) and with ATU (EMD-83349), have been deposited in the Electron Microscopy Data Bank (http://www.ebi.ac.uk/pdbe/emdb/). The corresponding atomic coordinates (PDB: 45KL and 45LB, respectively) have been deposited in the Protein Data Bank (http://www.rcsb.org), respectively. All other data generated or analyzed during this study are included in this published article and its supplementary information files.

## Acknowledgements

We sincerely thank the staff at the cryo-EM center of Southern University of Science and Technology for their technical support on the Cryo-EM and High-Performance Computation platforms. We thank Michael Wagner (University of Vienna, Austria) for helpful discussions. Z.L. is an investigator of SUSTech Institute for Biological Electron Microscopy. This work was supported by the National Natural Science Foundation of China (32570211 to Z.L., 32500022 to X.Y., 22577066 to W.P., and 42573078/42371064 to P.H.) and the Science, Technology and Innovation Commission of Shenzhen Municipality (JCYJ20240813094922030 to Z.L. and JCYJ20250604144527036 to X.Y.), and Shandong Provincial Natural Science Foundation (ZR2025QB36 to W.P.)

## Author Contributions

Z.L., X.Y., W.P. and P.H. supervised the project; T.M. and R.W. carried out cell culture; T.M. carried out membrane extraction and protein purification; T.M. and R.W. performed the biochemical assays; J.Y. and Z.L. prepared the samples for EM and collected the cryo-EM data; X.Y. analyzed cryo-EM data of *Ni*AMO and ATU-treated *Ni*AMO; X.Y., Z.L. and J.Y. performed the model building; J.Y., Z.L. and X.Y. conducted structural analysis; T.M. and R.W. conducted multi-omics experiments. Z.H. and W.P. designed and performed multiscale computational simulations; J.Y., X.Y., W.P. and Z.L. jointly wrote the initial draft; Z.L. W.P., P.H. and X.Y. edited the manuscript.

