## Supplementary information for "Catalytic and inhibitory architecture of comammox ammonia monooxygenase"

This file including:
Supplementary Figures 1-22
Supplementary Tables 1-6

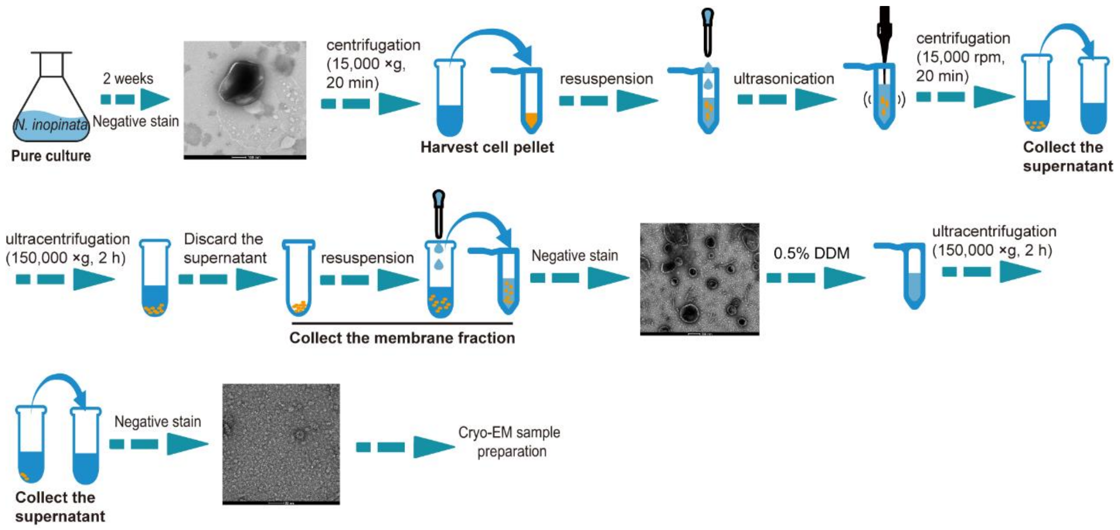

**Supplementary Fig. 1 | Schematic diagram depicting the purification of AMO from *N. inopinata*.** *Ni*AMO was purified in 0.5% DDM for cryo-EM sample preparation after checking by negative staining.

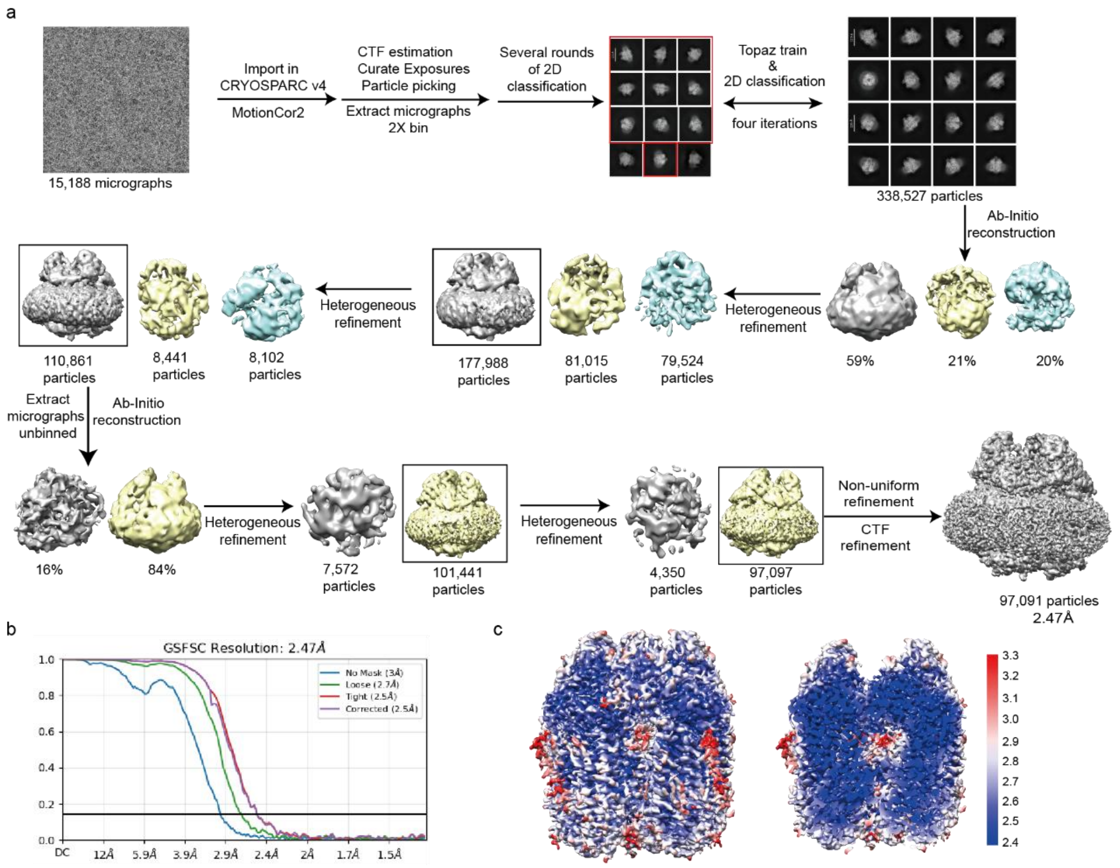

**Supplementary Fig. 2 | Cryo-EM data analysis of *Ni*AMO. a,** The flowchart of cryoEM data processing of *Ni*AMO. Details can be found in the "Method" section. **b,** Gold-standard Fourier shell correlation (GSFSC) curve for the *Ni*AMO map generated using cryoSPARC 4.3.2. The threshold of 0.143 was used to determine the overall resolution of the map. **c,** Local resolution for the overall reconstruction (left) and a central slice (right) of *Ni*AMO.

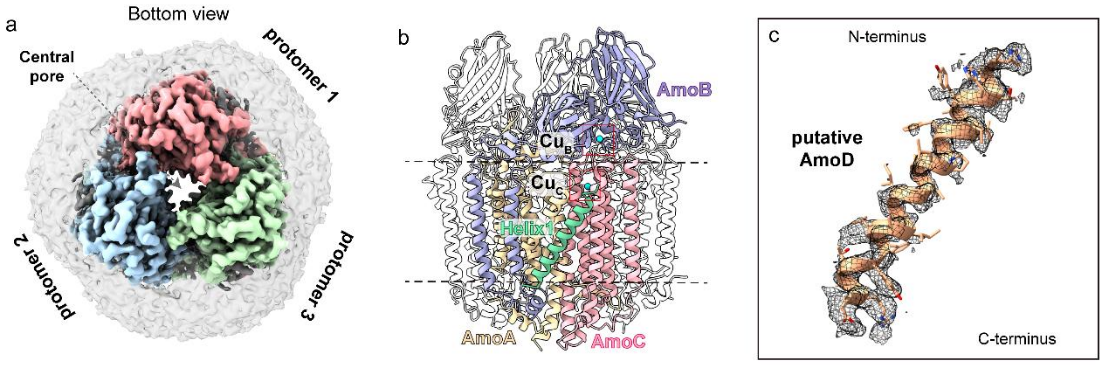

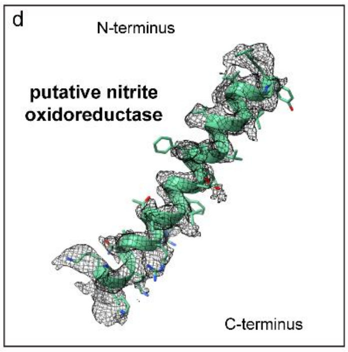

e

Mass-spectrometry analysis of *N. inopinata*

| Description | Score | Coverage % | # Unique Peptides | # Peptides | # PSM |
|---|---|---|---|---|---|
| AmoB | 125.298 | 46 | 16 | 16 | 189 |
| Putative nitrite oxidoreductase | 120.244 | 46 | 23 | 23 | 37 |
| SecD | 71.851 | 33 | 19 | 19 | 43 |
| Cytochrome c domain-containing protein | 52.906 | 25 | 9 | 9 | 22 |
| AmoA | 39.748 | 27 | 7 | 7 | 35 |
| SecF | 37.249 | 25 | 8 | 8 | 35 |
| AmoC | 20.955 | 16 | 3 | 3 | 5 |
| Putative AmoD1 | 18.889 | 22 | 3 | 3 | 7 |
| Putative AmoD2 | 7.626 | 14 | 3 | 3 | 5 |

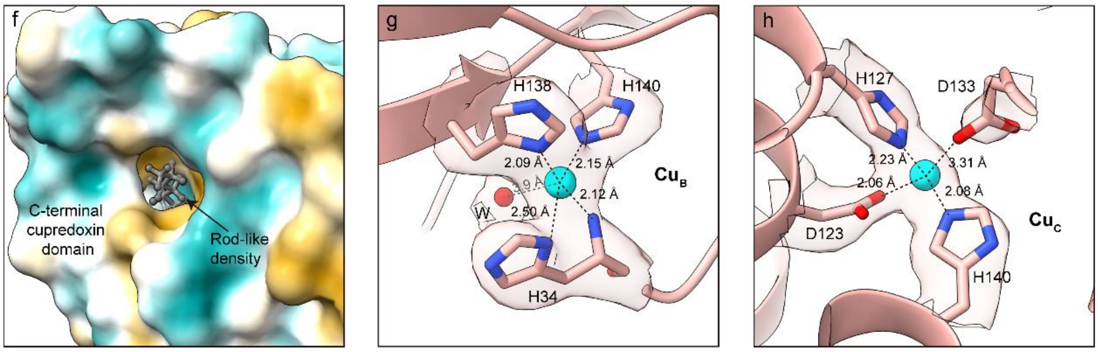

**Supplementary Fig. 3 | Cryo-EM structure of *Ni*AMO. a,** Cryo-EM structure from bottom view showing three protomers (colored in pink, blue and green, respectively) of *Ni*AMO. **b** Atomic model of *Ni*AMO monomer. Four components of *Ni*AMO are colored in yellow (AmoA), purple (AmoB), coral (AmoC) and green (Helix1). Cu$_B$ site and Cu$_C$ site are highlighted by red box, with their coppers colored in cyan. **c-d,** Helix1 with transparent cryo-EM density superimposed. **e,** Mass-spectrometric results of *N. inopinata* identifying peptide candidates for Helix1. Two of these candidates, highlighted by red boxes, were considered as potential matches. **f,** Rod-like density (colored in gray) located in the C-terminal cupredoxin domain of AmoB. The hydrophobic and hydrophilic surface characteristics of the cupredoxin domain are depicted. **g-h,** Cu$_B$ site (**e**) and Cu$_C$ site (**f**) and their coordination environments with transparent cryo-EM density superimposed.

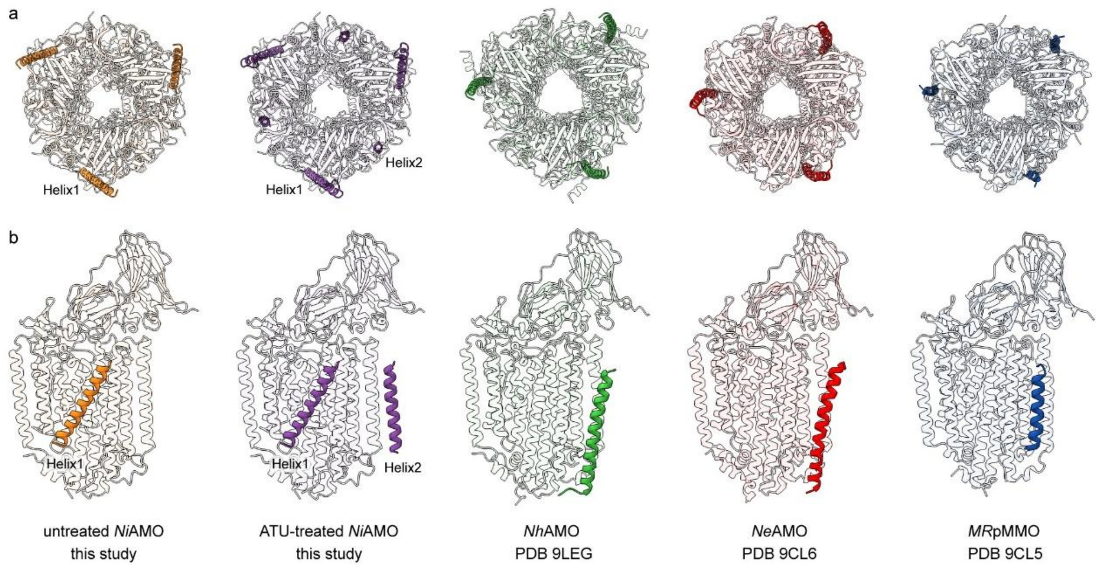

**Supplementary Fig. 4 | Comparison of AMO and pMMO determined by cryo-EM SPA.** The extra helices are highlighted in the corresponding atomic models of the monomer from top view (**a**) and the protomer from side view (**b**) of AMOs and pMMO. *Ni*AMO, ATU-treated NiAMO in this study, *Nh*AMO (PDB: 9LEG), *Ne*AMO (PDB: 9CL6) and *MR*pMMO (PDB: 9CL5) were colored in orange, purple, green, red and blue.

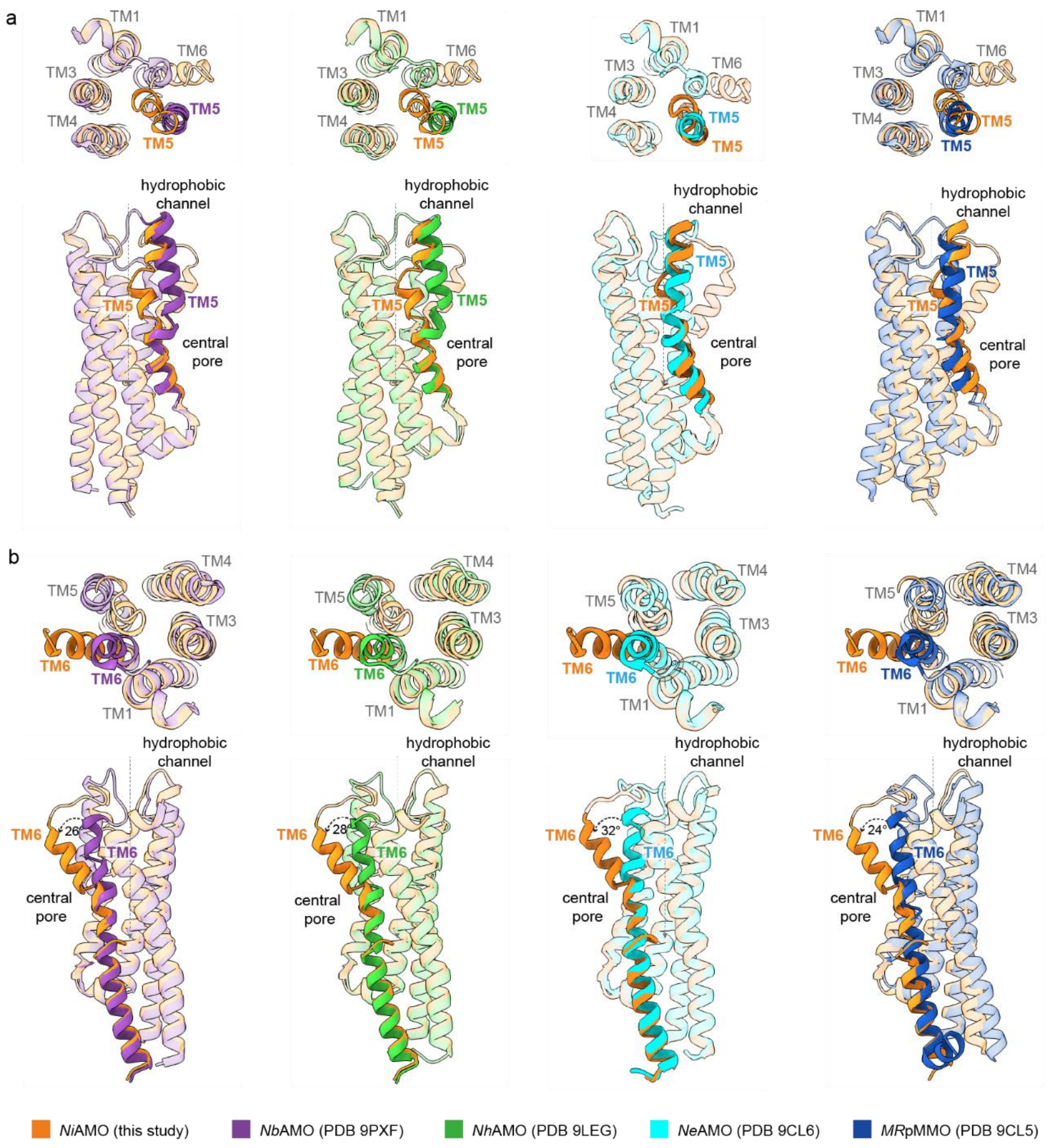

**Supplementary Fig. 5 | Comparison of hydrophobic channel from AMOs**. **a,** Alignment of conserved hydrophobic channel highlighting TM5 in Amoc from AMOs shown in top view (top) and side view (bottom). **b,** Alignment of conserved hydrophobic channel highlighting TM6 in Amoc from AMOs shown in top view (top) and side view (bottom). *Ni*AMO in this study, *Nb*AMO (PDB: 9PXF), *Nh*AMO (PDB: 9LEG), *Ne*AMO (PDB: 9CL6) and *MR*pMMO (PDB: 9CL5) were colored in orange, purple, green, cyan and blue.

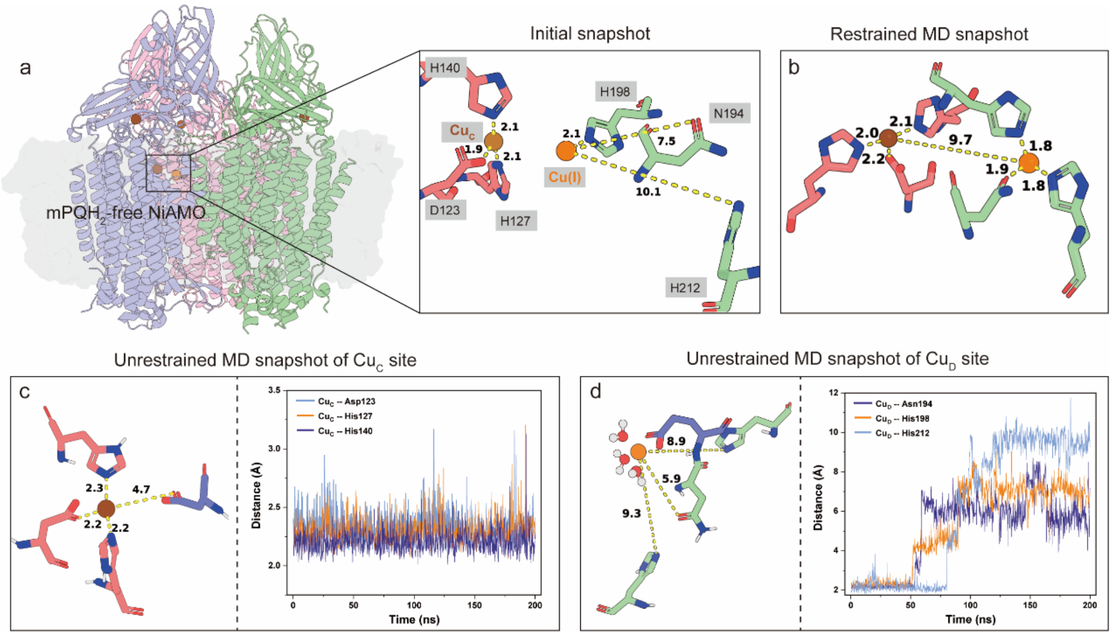

**Supplementary Fig. 6 | Evaluation of the Cu ion capture capability of the Cu$_D$ site in the mPQH$_2$-free *Ni*AMO. a,** Overall structure of membrane-bound mPQH$_2$-free *Ni*AMO and the initial configuration for MD simulations, in which a Cu(I) ion was randomly positioned near the Cu$_D$ site for conformational sampling. **b,** Stable Cu(I) coordination geometry at the Cu$_D$ site obtained from restrained MD simulations by applying distance and angle restraints between the Cu(I) ion and the surrounding coordinating residues. **c-d,** Representative conformations of the Cu$_C$ and Cu$_D$ sites obtained from unrestrained MD simulations without predefined coordination bonds, together with the time-dependent distances between the Cu(I) ions and their surrounding coordinating residues. The membrane is shown as a gray surface. The Cu(I) ions resolved in the cryo-EM structure are represented as brown spheres, whereas the Cu(I) ions randomly placed near the Cu$_D$ site are shown as orange spheres. The coordinating residues at the Cu$_C$ and Cu$_D$ sites are depicted as red and green sticks, respectively. Key distances are given in angstrom (Å).

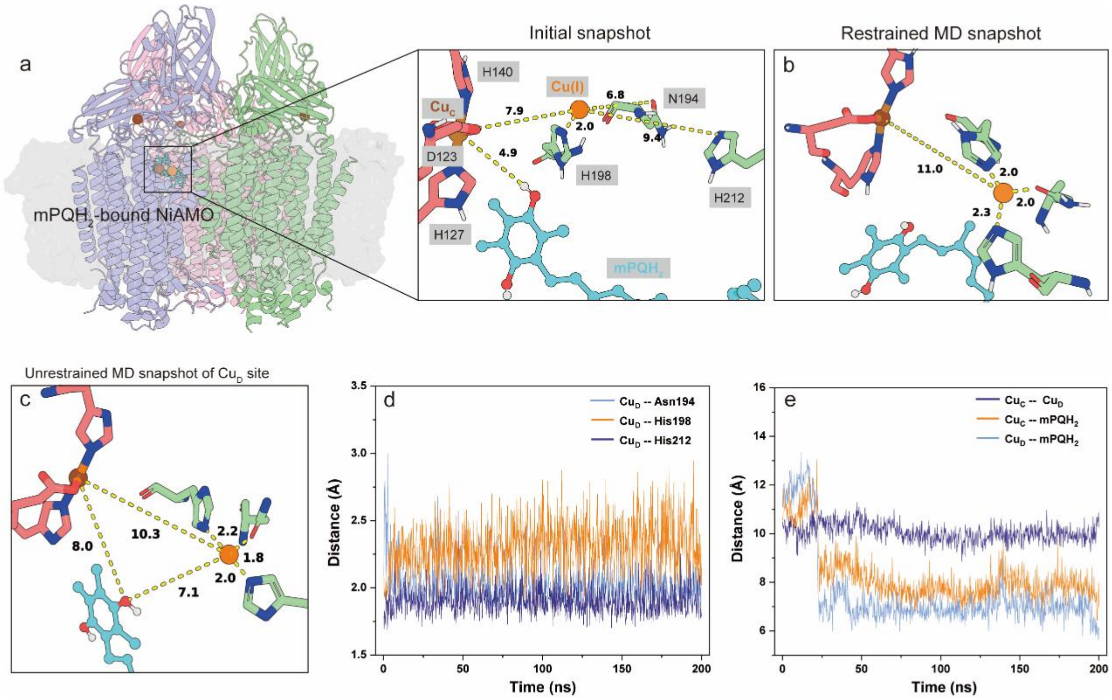

**Supplementary Fig. 7 | Evaluation of the Cu ion capture capability of the $Cu_D$ site in the $mPQH_2$-bound *Ni*AMO. a,** Overall structure of the membrane-bound *Ni*AMO-$mPQH_2$ complex and the initial snapshot for MD simulations, in which a Cu(I) ion was randomly positioned near the $Cu_D$ site for conformational sampling. **b,** Stable Cu(I) coordination geometry at the $Cu_D$ site obtained from restrained MD simulations by applying distance and angle restraints between the Cu(I) ion and the surrounding coordinating residues. **c,** Representative conformations from unrestrained MD simulations with coordination bond parameters applied only to the $Cu_C$ site. **d-e,** Time-dependent distances between the Cu(I) ion at the $Cu_D$ site and its surrounding coordinating residues, as well as $mPQH_2$. The membrane is shown as a gray surface. The Cu(I) ions and $mPQH_2$ resolved in the cryo-EM structure are represented as brown and cyan spheres, respectively, whereas the Cu(I) ions of $Cu_D$ site are shown as orange spheres. The coordinating residues at the $Cu_C$ and $Cu_D$ sites are depicted as red and green sticks, respectively. Key distances are given in angstrom (Å).

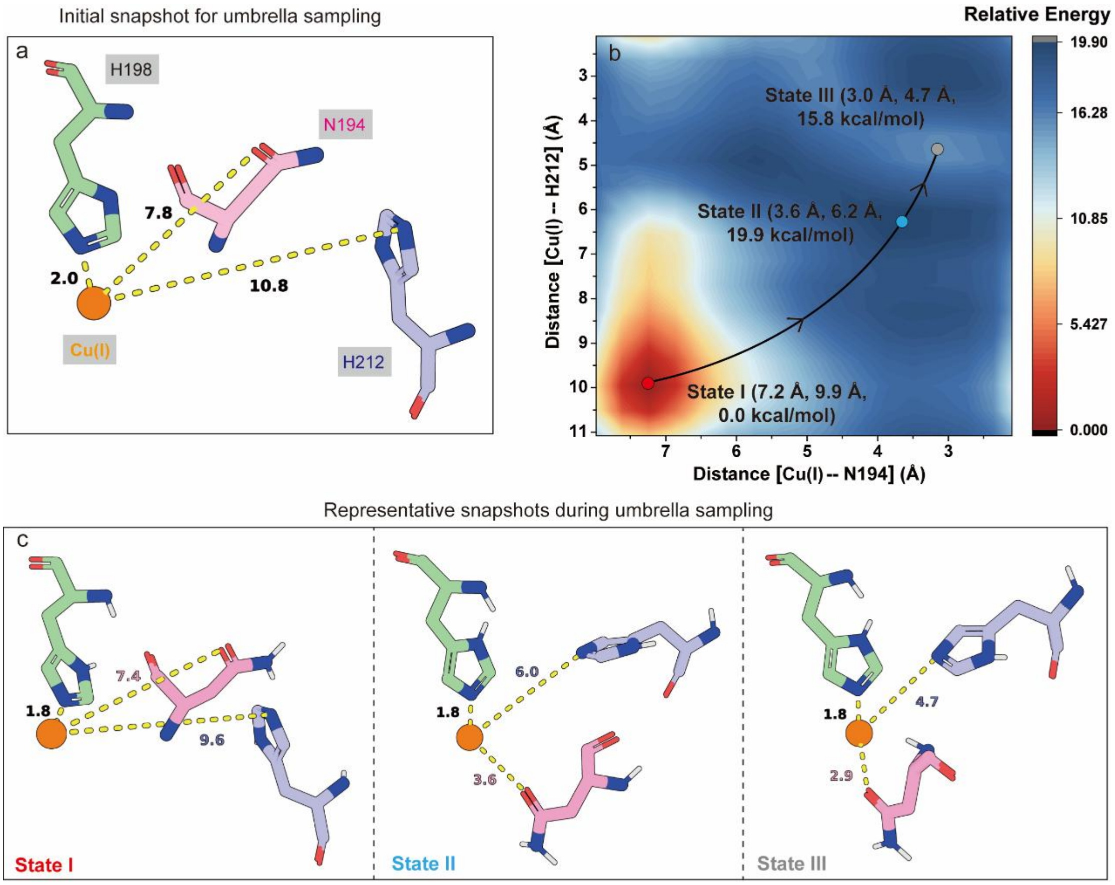

**Supplementary Fig. 8 | Free energy profile for Cu ion capture by the CuD site in the mPQH$_2$-free structure. a,** Initial snapshot for umbrella sampling simulations, in which Cu(I) was placed near H198. Asn194, His198, and His212 are depicted as pink, green, and blue stick models, respectively, while Cu(I) is represented as an orange sphere. **b,** Umbrella-sampling-calculated the free energy profile (in kcal/mol) for the transition of Cu(I) from the initial non-coordinating state (State I) to the Cu-coordinating state (State III). The reaction coordinates were defined as the distances between Cu(I) and N194 and between Cu(I) and H212, respectively. **c,** Representative snapshots sampled along the transition pathway of Cu(I) from the uncoordinated to the coordinated state. Key distances are given in angstrom (Å).

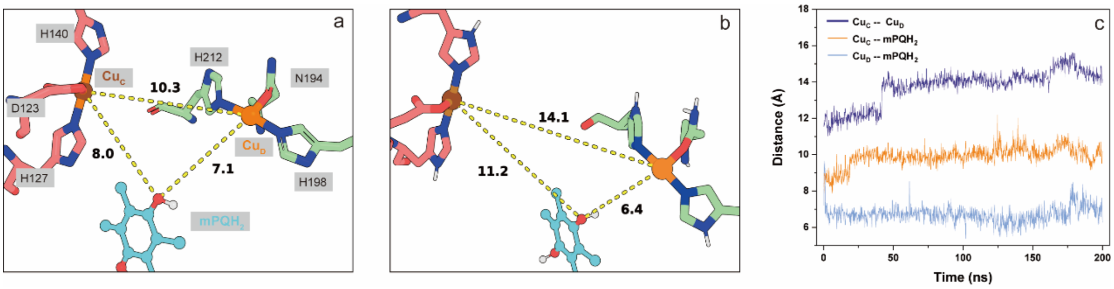

**Supplementary Fig. 9 | Binding mode in the active site of $mPQH_2$-bound *Ni*AMO. a-b,** The initial and representative configurations of *Ni*AMO containing $mPQH_2$ and simultaneously occupied $Cu_C$ and $Cu_D$ sites during MD simulations. **c,** Time-dependent distances between $mPQH_2$ and the $Cu_C$ and $Cu_D$ sites, as well as between the $Cu_C$ and $Cu_D$ copper ions. The $Cu_C$ and $Cu_D$ sites are shown as brown and orange spheres, respectively, whereas $mPQH_2$ is shown as cyan spheres. The coordinating residues at the $Cu_C$ and $Cu_D$ sites are depicted as red and green sticks, respectively. Key distances are given in angstrom (Å).

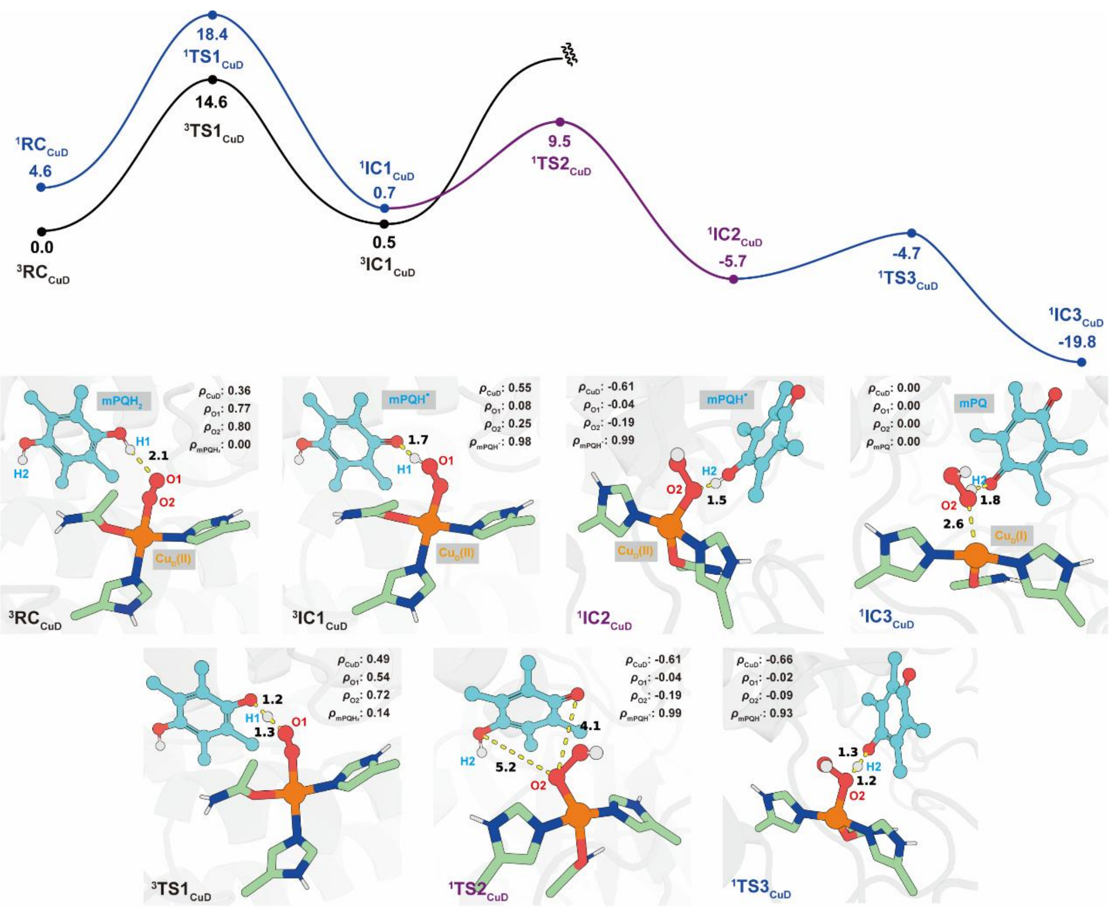

**Supplementary Fig. 10 | Calculated the mechanism of $Cu_D(I)\cdots H_2O_2$ intermediate generation.** QM(UB3LYP-D3/def2-TZVP)/MM relative energies (in kcal/mol) for $Cu_D(I)\cdots H_2O_2$ ($IC3_{CuD}$) formation from $Cu_D(II)–O2^{\bullet-}$ and $mPQH_2$ complex ($RC_{CuD}$) via the two HAT processes in the open-shell broken-symmetry singlet and triplet states. QM(UB3LYP-D3/def2-SVP)/MM optimized geometries of key species involved in the reaction are presented. Key distances are given in angstrom (Å).

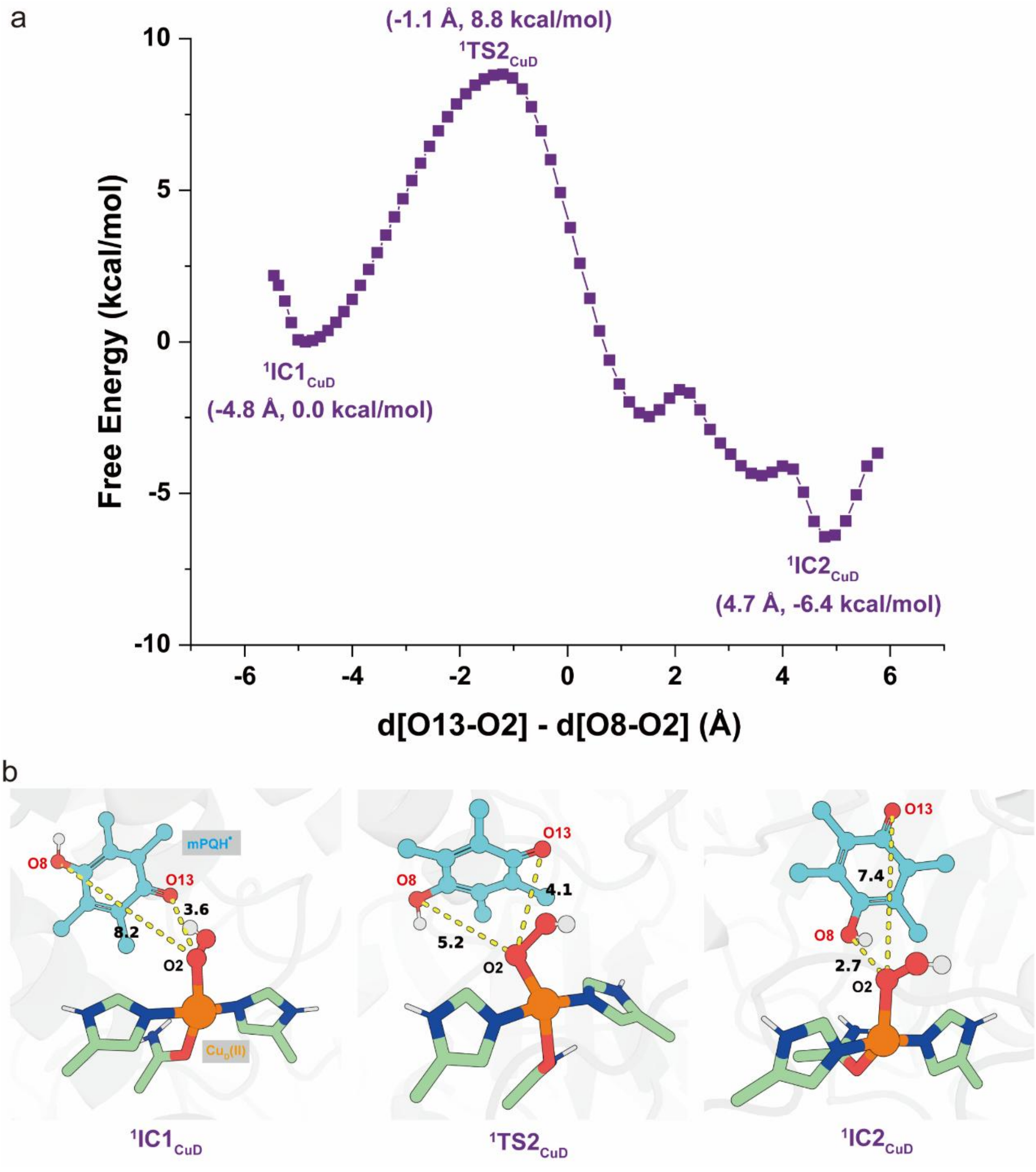

**Supplementary Fig. 11 | Calculated the mechanism of mPQH$^{\bullet}$ radical conformational reorganization near the Cu$_D$ site. a,** QM/MM metadynamics-calculated free energy profile (in kcal/mol) for the conformational reorganization of the mPQH$^{\bullet}$ radical near the Cu$_D$ site. The reaction coordinate is defined as the difference between the distance from O2 of the Cu$_D$(II)−OOH$^{-}$ species to O13 of the mPQH$^{\bullet}$ radical (d1) and the distance between O2 and O8 of the mPQH$^{\bullet}$ radical (d2). **b,** The geometry of intermediates and transition states. Key distances are given in angstrom (Å).

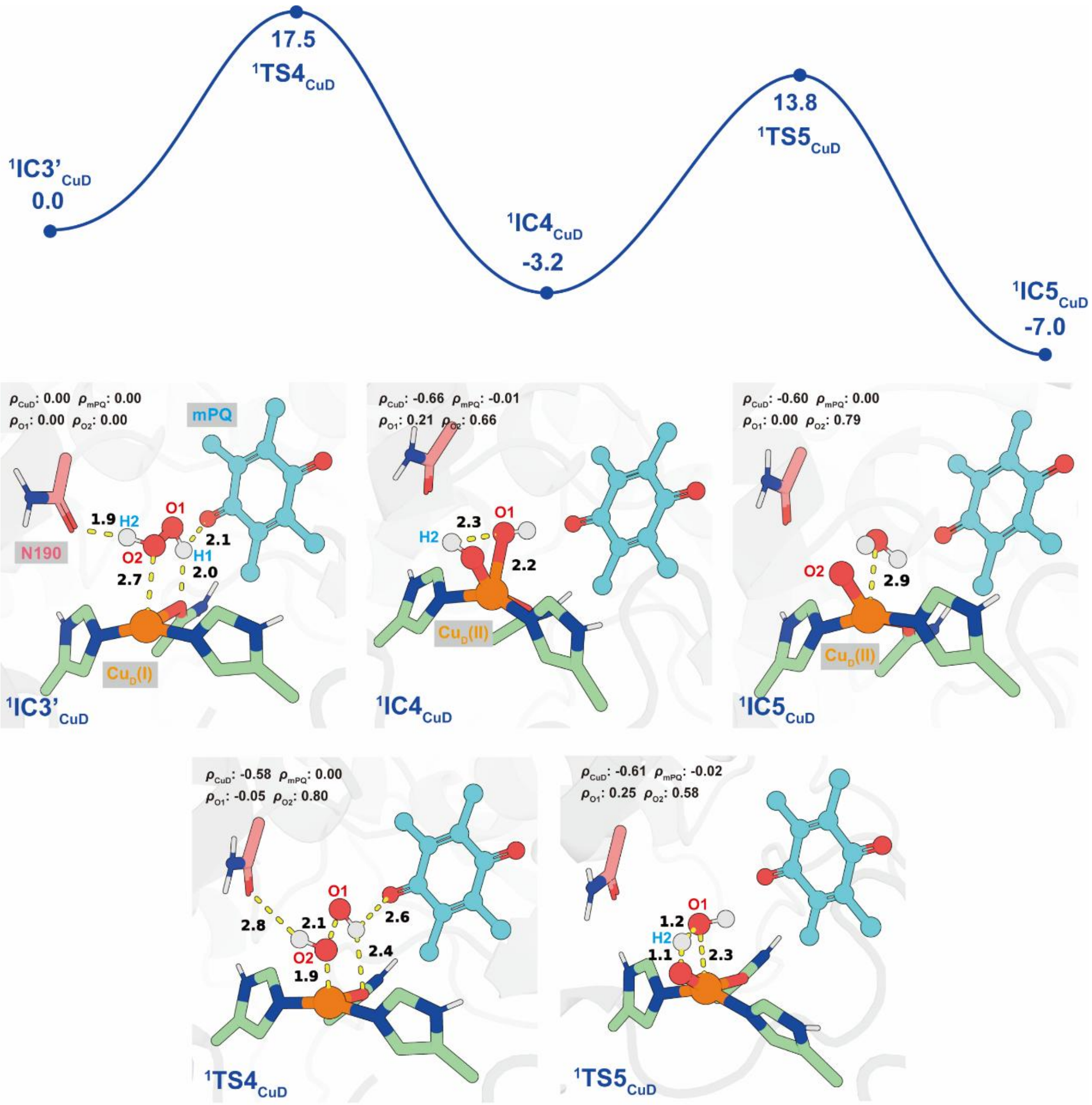

**Supplementary Fig. 12 | Calculated the mechanism of reactive oxygen species generation.** QM(UB3LYP-D3/def2-TZVP)/MM relative energies (in kcal/mol) for $Cu_D(II){-}O^{\bullet-}$ ($^1IC5_{CuD}$) formation from $Cu_D(I)\cdots H_2O_2$ ($^1IC3'_{CuD}$) in the open-shell broken-symmetry singlet state. QM(UB3LYP-D3/def2-SVP)/MM optimized geometries of key species involved in the reaction are presented. Key distances are given in angstrom (Å).

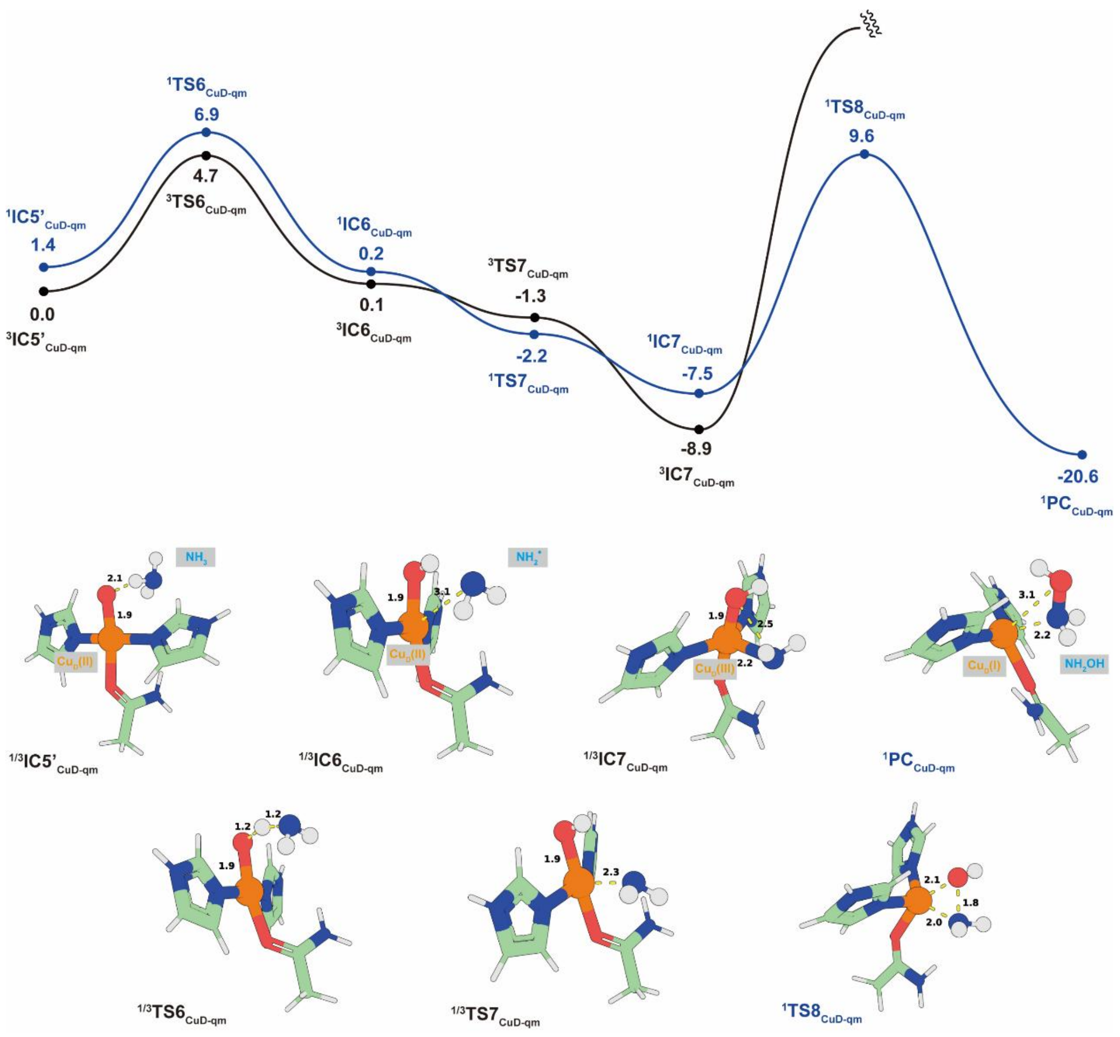

**Supplementary Fig. 13 | Calculated mechanism of Cu$_D$-mediated ammonia oxidation.** QM (UB3LYP/def2-TZVP//def2-SVP) calculated potential energy profile (in kcal/mol) for Cu$_D$(II)−O$^{\bullet-}$ mediated ammonia hydroxylation to hydroxylamine via the HAT/oxygen rebound mechanism in both open-shell broken-symmetry singlet and triplet states. Key distances are given in angstrom (Å). The spin density population of key atoms for all species are summarized in Supplementary Table S3.

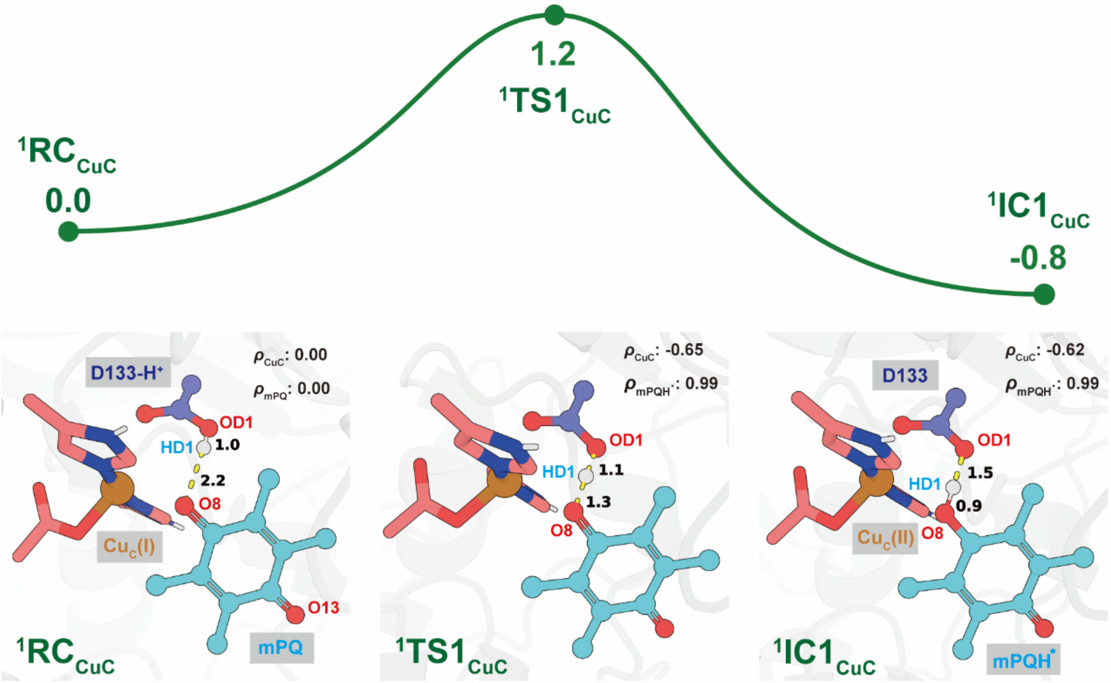

**Supplementary Fig. 14 | Calculated mechanism of the first proton-coupled reduction of mPQ to the mPQH$^{\bullet}$ radical at the Cu$_C$ site.** QM(UB3LYP-D3/def2-TZVP)/MM-calculated relative energies (in kcal/mol) for the first proton-coupled electron transfer (PCET) from oxidized mPQ ($^1$RC$_{CuC}$) to form the mPQH$^{\bullet}$ radical ($^1$IC1$_{CuC}$) at the Cu$_C$ site in the open-shell broken-symmetry singlet state. QM(UB3LYP-D3/def2-SVP)/MM optimized geometries of key species involved in the reaction are presented. The proton donor residue D133 is highlighted in a blue ball-and-stick model. Key distances are given in angstrom (Å).

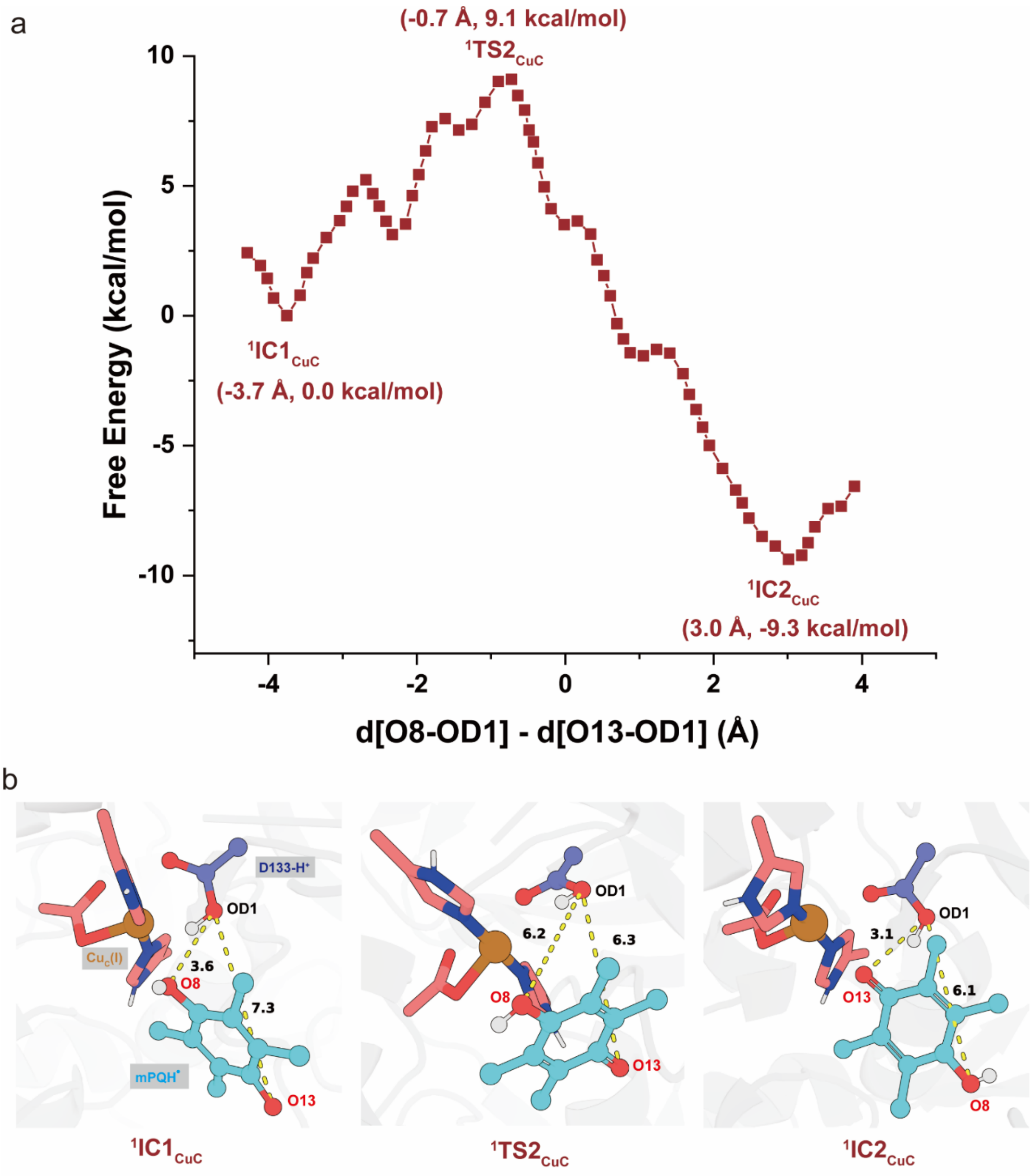

**Supplementary Fig. 15 | Calculated the mechanism of mPQH• radical conformational reorganization near the Cu$_C$ site. a,** QM/MM metadynamics-calculated free energy profile (in kcal/mol) for the conformational reorganization of the mPQH• radical near the Cu$_C$ site. The reaction coordinate is defined as the difference between the distance from OD1 of the residue D133 to O8 of the mPQH• radical (d1) and the distance between OD1 and O13 of the mPQH• radical (d2). **b,** The geometry of intermediates and transition states. Key distances are given in angstrom (Å).

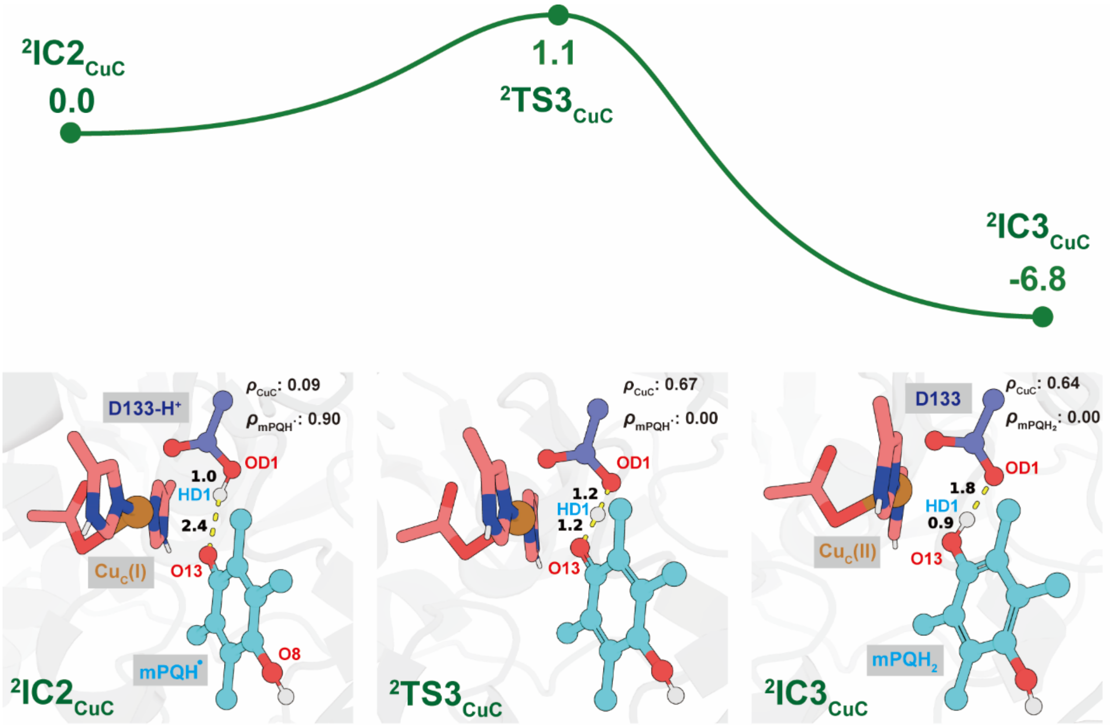

**Supplementary Fig. 16 | Calculated mechanism of the second proton-coupled reduction of the $mPQH^{\bullet}$ radical to $mPQH_2$ at the $Cu_C$ site.** QM(UB3LYP-D3/def2-TZVP)/MM-calculated relative energies (in kcal/mol) for the second proton-coupled electron transfer (PCET) from the $mPQH^{\bullet}$ radical ($^2IC2_{CuC}$) to form $mPQH_2$ ($^2IC3_{CuC}$) at the $Cu_C$ site in the open-shell double state. QM(UB3LYP-D3/def2-SVP)/MM-optimized geometries of key species involved in the reaction are presented. The proton donor residue D133 is highlighted in a blue ball-and-stick model. Key distances are given in angstroms (Å).

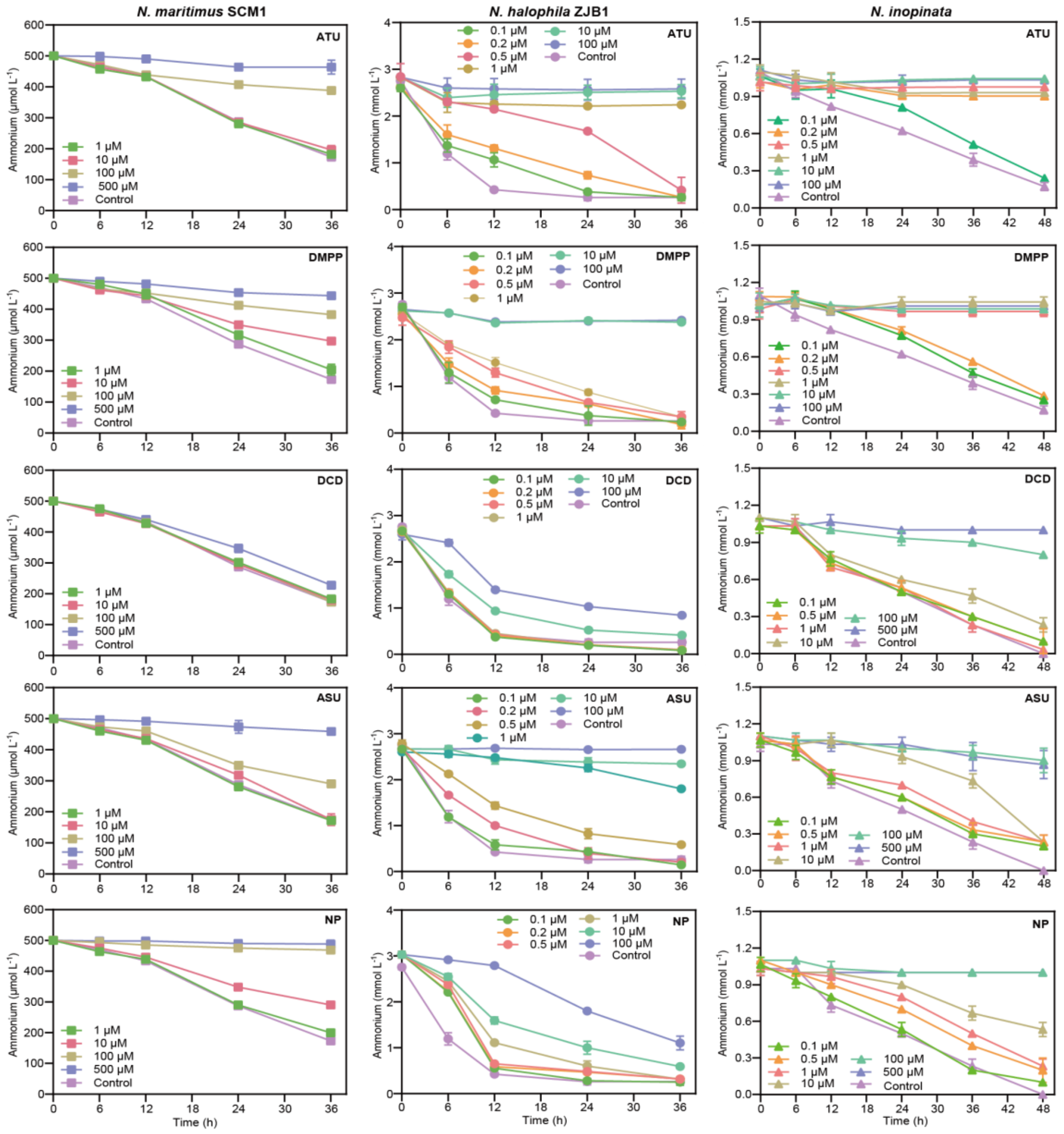

**Supplementary Fig. 17 | Average inhibition of AO in AOA (left panels), AOB (middle panels) and comammox (right panels) under varying concentrations of NIs.** Each subplot corresponds to a specific NI: ATU, DMPP, DCD, ASU and NP. Different colors and symbols represent different concentrations of the respective inhibitor. Control treatments lack NIs. Error bars indicate standard deviations derived from ⩾3 biological replicates, each performed in triplicate. These data correspond to Fig. 3b.

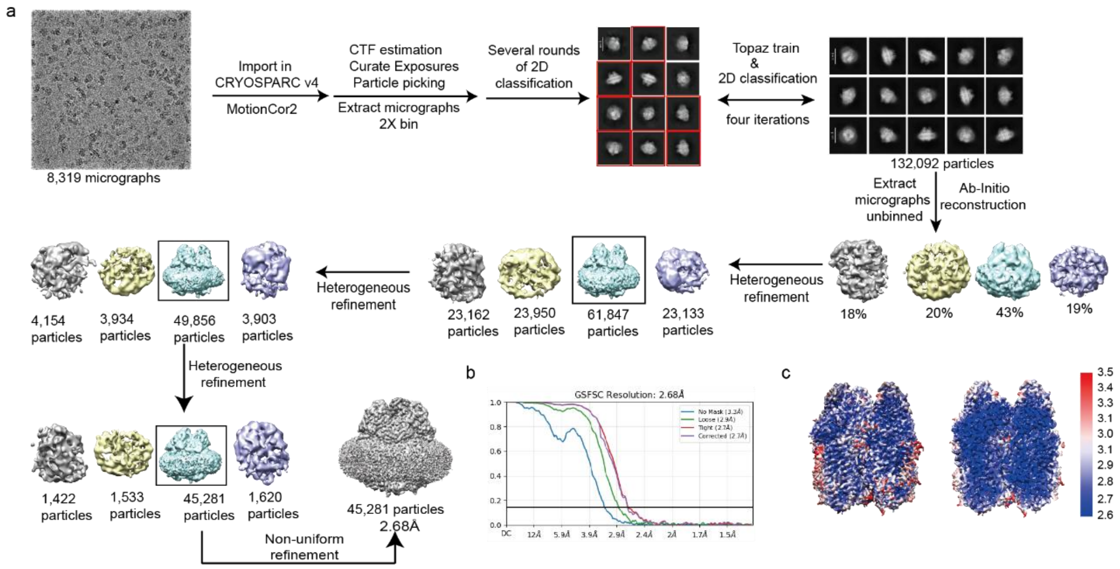

**Supplementary Fig. 18 | Cryo-EM data analysis of ATU-treated *Ni*AMO. a,** The flowchart of cryoEM data processing of ATU-treated *Ni*AMO. Details can be found in the “Method” section. **b,** Gold-standard Fourier shell correlation (GSFSC) curve for the ATU-treated *Ni*AMO map generated using cryoSPARC 4.3.2. The threshold of 0.143 was used to determine the overall resolution of the map. **c,** Local resolution for the overall reconstruction (left) and a central slice (right) of ATU-treated *Ni*AMO.

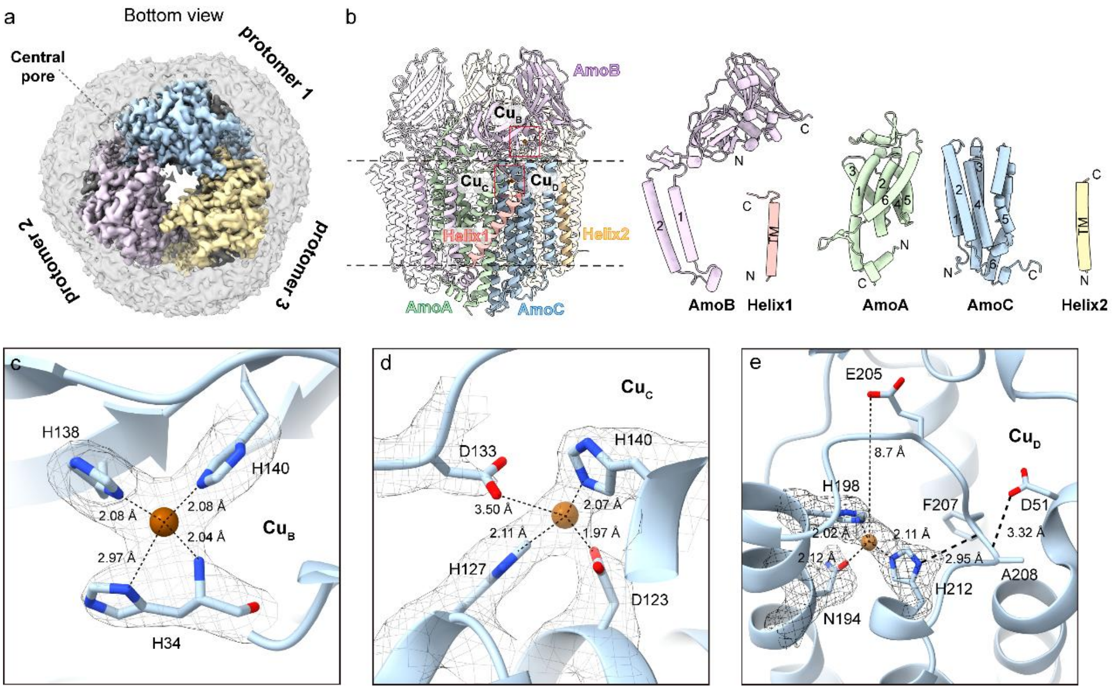

**Supplementary Fig. 19 | Cryo-EM structure of ATU-treated *Ni*AMO. a,** Cryo-EM structure from bottom view showing three protomers (colored in blue, light purple and yellow, respectively) of ATU-treated *Ni*AMO. **b,** Atomic model of ATU-treated *Ni*AMO monomer. Five components of ATU-treated *Ni*AMO are colored in green (AmoA), purple (AmoB), blue (AmoC), coral (Helix1) and yellow (Helix2). $Cu_B$ site, $Cu_C$ site and $Cu_D$ site are highlighted by red box, with their coppers colored in brown. **c-e,** $Cu_B$ site (**c**), $Cu_C$ site (**d**) and $Cu_D$ site (**e**) and their coordination environments with mesh cryo-EM density superimposed.

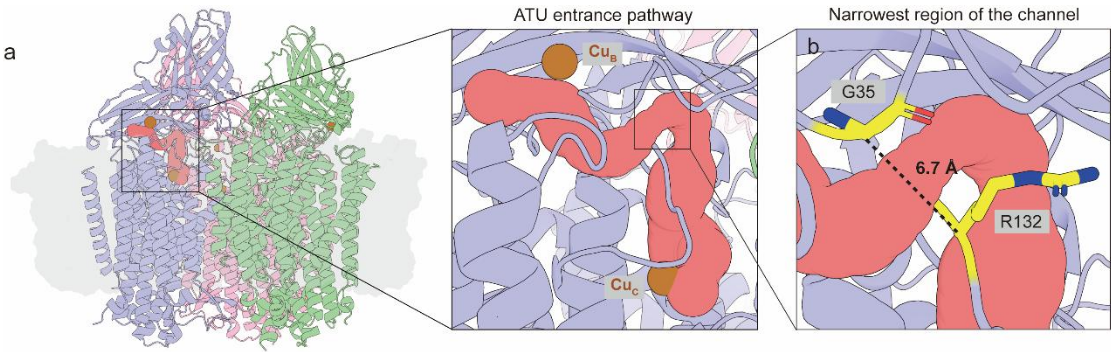

**Supplementary Fig. 20 | Pathway of ATU migration into the *Ni*AMO active site. a,** Overview of the channel pathway mediating ATU transport from the external environment to the active site of *Ni*AMO. **b,** The narrowest region of the channel for ATU entry. The amino acid residues at the narrowest regions are displayed as yellow stick models.

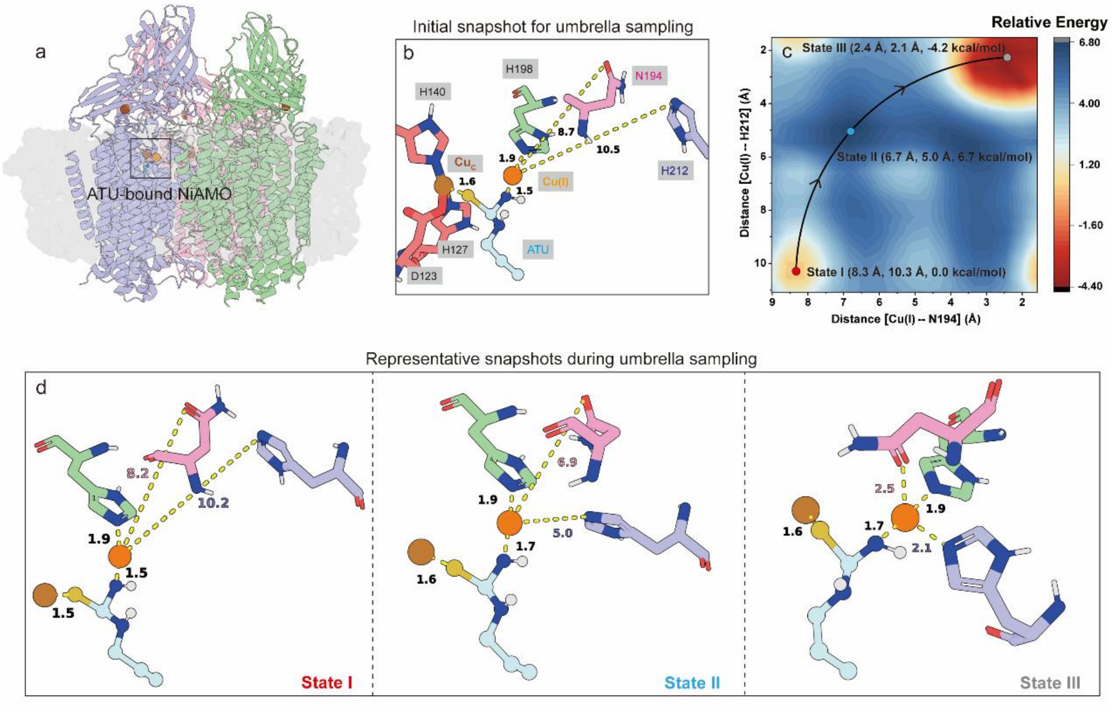

**Supplementary Fig. 21 | Free-energy profile for Cu(I) ion capture by the $Cu_D$ site in *Ni*AMO following ATU entry. a,** Overall structure of apo *Ni*AMO with ATU and the Cu(I) ion at the $Cu_D$ site superimposed based on the cryo-EM structure of ATU-treated *Ni*AMO. The membrane is shown as a gray surface. Cu(I) ions at the $Cu_C$ and $Cu_D$ sites are shown as brown and orange spheres, respectively, and ATU is shown as a cyan ball-and-stick model. **b,** Initial snapshot for umbrella sampling simulations, with ATU and the Cu(I) ion at the $Cu_D$ site positioned according to the ATU-treated *Ni*AMO structure. $Cu_C$-site coordinating residues are shown as red sticks, while $Cu_D$-site residues N194, H198, and H212 are shown in pink, green, and blue, respectively. **c,** Umbrella-sampling-calculated the free energy profile (in kcal/mol) for the transition of Cu(I) from the initial non-coordinating state (State I) to the Cu-coordinating state (State III). The reaction coordinates were defined as the distances between Cu(I) and N194 and between Cu(I) and H212, respectively. **d,** Representative snapshots sampled along the transition pathway of Cu(I) from the uncoordinated to the coordinated state. Key distances are given in angstrom (Å).

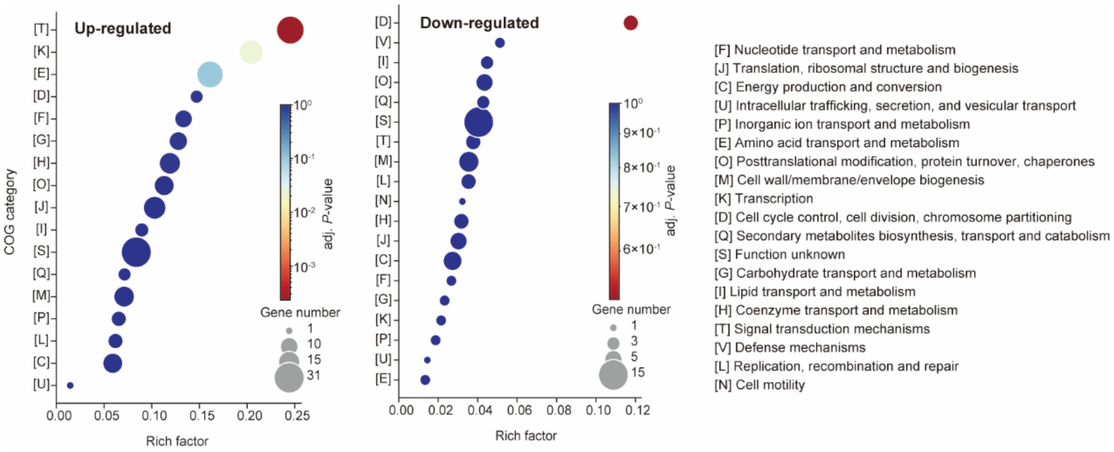

**Supplementary Fig. 22 | Enrichment analysis of differentially proteins based on EggNOG database in every cluster from proteomic heatmap.** Thresholds: $|\log_2 FC| > 1.2$ and adjusted $P < 0.05$.

**Supplementary Table 1. QM/MM energies (in Hartree, a.u.) of all species**

| | UB3LYP/def2-TZVP//def2-SVP | | | |
|---|---|---|---|---|
| **Species** | **QM** | **MM** | **QM/MM** | **ZPE** |
| $^{1}RC_{CuD}$ | -3221.5200 | -90.0915 | -3311.6115 | 0.5520 |
| $^{3}RC_{CuD}$ | -3221.5261 | -90.0928 | -3311.6189 | 0.5057 |
| $^{1}TS1_{CuD}$ | -3221.5007 | -90.0867 | -3311.5874 | 0.5498 |
| $^{3}TS1_{CuD}$ | -3221.5048 | -90.0884 | -3311.5932 | 0.5032 |
| $^{1}IC1_{CuD}$ | -3221.5289 | -90.0887 | -3311.6176 | 0.5518 |
| $^{3}IC1_{CuD}$ | -3221.5282 | -90.0895 | -3311.6177 | 0.5053 |
| $^{1}IC2_{CuD}$ | -3221.4708 | -56.9803 | -3278.4511 | 0.5490 |
| $^{1}TS3_{CuD}$ | -3221.4696 | -56.9782 | -3278.4479 | 0.5473 |
| $^{1}IC3_{CuD}$ | -3221.4583 | -57.0139 | -3278.4722 | 0.5477 |
| $^{1}IC3'_{CuD}$ | -3430.6902 | -54.1615 | -3484.8517 | 0.6270 |
| $^{1}TS4_{CuD}$ | -3430.6571 | -54.1588 | -3484.8159 | 0.6191 |
| $^{1}IC4_{CuD}$ | -3430.6918 | -54.1612 | -3484.8530 | 0.6233 |
| $^{1}TS5_{CuD}$ | -3430.6608 | -54.1618 | -3484.8226 | 0.6200 |
| $^{1}IC5_{CuD}$ | -3430.7034 | -54.1558 | -3484.8593 | 0.6234 |
| $^{1}RC_{CuC}$ | -3167.8553 | -58.9868 | -3226.8421 | / |
| $^{1}TS1_{CuC}$ | -3167.8591 | -58.9812 | -3226.8403 | / |
| $^{1}IC1_{CuC}$ | -3167.8621 | -58.9814 | -3226.8435 | / |
| $^{2}IC2_{CuC}$ | -3168.5869 | -59.0611 | -3227.6480 | / |
| $^{2}TS2_{CuC}$ | -3168.5938 | -59.0525 | -3227.6462 | / |
| $^{2}IC3_{CuC}$ | -3168.6068 | -59.0520 | -3227.6588 | / |

**Supplementary Table 2. QM energies (in Hartree, a.u.) of all species**

| | UB3LYP/def2-TZVP//def2-SVP | | |
|---|---|---|---|
| **Species** | **EE** | **ZPE** | **EE+ZPE** |
| $^{1}$IC5’$_{CuD}$ | -2434.244 | 0.262 | -2433.981 |
| $^{1}$TS6$_{CuD}$ | -2434.229 | 0.256 | -2433.973 |
| $^{1}$IC6$_{CuD}$ | -2434.242 | 0.259 | -2433.983 |
| $^{1}$TS7$_{CuD}$ | -2434.243 | 0.258 | -2433.985 |
| $^{1}$IC7$_{CuD}$ | -2434.256 | 0.261 | -2433.996 |
| $^{1}$TS8$_{CuD}$ | -2434.228 | 0.260 | -2433.968 |
| $^{1}$PC$_{CuD}$ | -2434.281 | 0.264 | -2434.016 |
| $^{3}$IC5’$_{CuD}$ | -2434.246 | 0.262 | -2433.984 |
| $^{3}$TS6$_{CuD}$ | -2434.232 | 0.256 | -2433.976 |
| $^{3}$IC6$_{CuD}$ | -2434.242 | 0.259 | -2433.984 |
| $^{3}$TS7$_{CuD}$ | -2434.244 | 0.258 | -2433.986 |
| $^{3}$IC7$_{CuD}$ | -2434.259 | 0.261 | -2433.999 |

**Supplementary Table 3. Spin density population of key atoms for all species**

| Species | Spin density | | | |
|---|---|---|---|---|
| **QM/MM model** | **$Cu_D$** | **O1** | **O2** | **$mPQH_2$/ $mPQH^{\bullet}$ / mPQ** |
| $^{3}RC_{CuD}$ | 0.36 | 0.77 | 0.8 | 0 |
| $^{3}TS1_{CuD}$ | 0.49 | 0.54 | 0.72 | 0.14 |
| $^{3}IC1_{CuD}$ | 0.55 | 0.08 | 0.25 | 0.98 |
| $^{1}RC_{CuD}$ | -0.45 | 0.37 | 0.18 | 0 |
| $^{1}TS1_{CuD}$ | -0.57 | 0.37 | 0.26 | 0.09 |
| $^{1}IC1_{CuD}$ | -0.56 | -0.07 | -0.26 | 0.99 |
| $^{1}TS2_{CuD}$ | -0.61 | -0.04 | -0.19 | 0.99 |
| $^{1}IC2_{CuD}$ | -0.61 | -0.04 | -0.19 | 0.99 |
| $^{1}TS3_{CuD}$ | -0.66 | -0.02 | -0.09 | 0.93 |
| $^{1}IC3_{CuD}$ | 0 | 0 | 0 | 0 |
| $^{1}IC3'_{CuD}$ | 0 | 0 | 0 | 0 |
| $^{1}TS4_{CuD}$ | -0.58 | 0.8 | -0.05 | 0 |
| $^{1}IC4_{CuD}$ | -0.66 | 0.66 | 0.21 | -0.01 |
| $^{1}TS5_{CuD}$ | -0.61 | 0.58 | 0.25 | -0.02 |
| $^{1}IC5_{CuD}$ | -0.6 | 0.79 | 0 | 0 |
| **QM model** | **$Cu_D$** | **O** | | **$NH_3$/ $NH_2^{\bullet}$/ NH2OH** |
| $^{1}IC5'_{CuD}$ | -0.62 | 0.83 | | 0.0 |
| $^{1}TS6_{CuD}$ | -0.58 | 0.32 | | 0.52 |
| $^{1}IC6_{CuD}$ | -0.59 | -0.06 | | 0.90 |
| $^{1}TS7_{CuD}$ | -0.57 | -0.24 | | 0.97 |
| $^{1}IC7_{CuD}$ | -0.52 | 0.08 | | 0.57 |

| | | | |
|---|---|---|---|
| $^{1}$TS8$_{CuD}$ | -0.28 | 0.05 | 0.27 |
| $^{1}$PC$_{CuD}$ | 0.0 | 0.0 | 0.0 |
| $^{3}$IC5'$_{CuD}$ | 0.50 | 1.24 | 0.0 |
| $^{3}$TS6$_{CuD}$ | 0.54 | 0.73 | 0.56 |
| $^{3}$IC6$_{CuD}$ | 0.59 | 0.31 | 0.91 |
| $^{3}$TS7$_{CuD}$ | 0.59 | 0.24 | 0.97 |
| $^{3}$IC7$_{CuD}$ | 0.59 | 0.30 | 0.85 |
| **QM/MM model** | **Cu$_{C}$** | **mPQH$_2$/ mPQH$^{\cdot}$/ mPQ** | |
| $^{1}$RC$_{CuC}$ | 0.0000 | 0.0000 | |
| $^{1}$TS1$_{CuC}$ | -0.6500 | 0.9900 | |
| $^{1}$IC1$_{CuC}$ | -0.6200 | 0.9900 | |
| $^{2}$IC2$_{CuC}$ | 0.0900 | 0.9000 | |
| $^{2}$TS3$_{CuC}$ | 0.6700 | 0.0000 | |
| $^{2}$IC3$_{CuC}$ | 0.6400 | 0.0000 | |

**Supplementary Table 4. Data collection, refinement and validation statistics**

| | PDB 45KL<br>EMDB-83338 | PDB 45LB<br>EMDB-83349 |
|---|---|---|
| **Data collection and processing** | | |
| EMDB code | 83338 | 83349 |
| PDB code | 45KL | 45LB |
| Magnification | 130,000 | 130,000 |
| Voltage (kV) | 300 | 300 |
| Camera | Falcon4i | Falcon4i |
| Electron exposure ($e^-$/Å$^2$) | 50 | 50 |
| Defocus range (µm) | -1.2 ~ -2.0 | -1.2 ~ -2.0 |
| Pixel size (Å) | 0.6584 | 0.6584 |
| Micrographs (no.) | 15,188 | 8,319 |
| Initial particle images (no.) | 338,527 | 132,092 |
| Final particle images (no.) | 97,091 | 45,281 |
| Symmetry imposed | C3 | C3 |
| Map resolution (Å) | 2.47 | 2.68 |
| Map sharpen B factor (Å$^2$) | 85.4 | 84.4 |
| FSC threshold | 0.143 | 0.143 |
| Map resolution range (Å) | 2.4-3.3 | 2.6-3.5 |
| **Refinement** | | |
| Model resolution (Å) | 2.47 | 2.68 |
| FSC threshold | 0.143 | 0.143 |
| **Model composition** | | |
| Non-hydrogen atoms | 23168 | 23555 |
| Protein residues | 2820 | 2898 |
| Ligands | CU: 6 | CU: 9, ATU: 3 |
| Waters | 476 | 437 |
| **ADP (B-factors)** | | |
| Protein | 8.54/125.85/52.45 | 3.46/141.00/40.10 |
| Ligand | 73.82/98.55/85.83 | 43.66/101.55/61.16 |
| Water | 30.87/75.25/47.89 | 13.96/71.87/38.02 |
| **R.m.s. deviations** | | |
| Bond lengths (Å) | 0.004 | 0.003 |
| Bond angles (°) | 0.645 | 0.554 |
| **Validation** | | |
| MolProbity score | 2.07 | 1.68 |
| Clashscore | 6.09 | 5.41 |
| Rotamer outliers (%) | 2.46 | 1.57 |
| **Ramachandran plot** | | |
| Favored (%) | 93.24 | 96.41 |
| Allowed (%) | 6.15 | 3.49 |
| Disallowed (%) | 0.61 | 0.10 |

**Supplementary Table 5. Copper content in active and inactivated *Ni*AMO samples measured by ICP-MS**

| Sample type | Sample name | Sample weight (g) | Final vol. (mL) | Element | Measured conc. (µg/L) | Cu (mg/kg)[a] |
|---|---|---|---|---|---|---|
| Active *Ni*AMO | AMO_WT-1 | 0.011 | 10 | Cu | 112.00 | 104.67[b] |
| | AMO_WT-2 | 0.011 | 10 | Cu | 111.80 | 104.48 |
| | AMO_WT-3 | 0.011 | 10 | Cu | 111.62 | 104.31 |
| Inactivated *Ni*AMO | AMO_ATU-1 | 0.0051 | 10 | Cu | 52.67 | 103.27 |
| | AMO_ATU-2 | 0.0051 | 10 | Cu | 52.67 | 103.28 |
| | AMO_ATU-3 | 0.0051 | 10 | Cu | 52.65 | 103.23 |
| Buffer | buffer-1 | 0.0049 | 10 | Cu | 2.21 | 4.50 |
| | buffer-2 | 0.0049 | 10 | Cu | 2.17 | 4.44 |
| | buffer-3 | 0.0049 | 10 | Cu | 2.25 | 4.59 |

[a]the net copper mass in the sample was calculated as: $m(Cu)_{net} = m(Cu)_{sample} - m(Cu)_{buffer}$. Analogous to pMMO, AMO accounted for approximately 80% of the total membrane-bound protein abundance.

[b]the molecular weight of *Nm*AMO trimer is 328.53 kDa.

**Supplementary Table 6. QPCR-based quantification of *amoA* gene abundance in AOA, AOB, and comammox**

| Target genes | Primer sequence (5′-3′) | PCR conditions | Efficiency | $R^2$ | References |
|---|---|---|---|---|---|
| AOA-*amoA* | 1F: STAATGGTCTGGCTTAGACG<br>2R: GCGGCCATCCATCTGTATGT | 95°C for 10min; 40× (95°C for 30s; 56°C for 45s; 72°C for 50s). | 92% | 0.99 | Francis et al., 2005[1] |
| *β*-AOB-*amoA* | 1F: GGGGHTTYTACTGGTGGT<br>2R: CCCCTCKGSAAAGCCTTCTTC | 95°C for 10min; 40× (95°C for 30s; 58°C for 40s; 72°C for 60s). | 85% | 0.99 | Kowalchuk et al., 1999[2] |
| CMX clade A-*amoA* | Ntsp-162F: GGATTTCTGGNTSGATTGGA<br>Ntsp-359R: WAGTTNGACCACCASTACCA | 95°C for 10min; 40× (95°C for 15s; 48°C for 30s; 72°C for 45s). | 85% | 0.99 | Fowler et al., 2018[3] |
